# Age of diabetes onset in the mutant proinsulin syndrome correlates with mutational impairment of protein foldability and stability

**DOI:** 10.1101/2021.09.24.461687

**Authors:** Balamurugan Dhayalan, Yen-Shan Chen, Yanwu Yang, Mark Jarosinski, Deepak Chatterjee, Rachel Grabowski, Shayan Poordian, Nelson F.B. Phillips, Peter Arvan, Faramarz Ismail-Beigi, Michael A. Weiss

## Abstract

Diverse heterozygous mutations in the human insulin gene cause a monogenic diabetes mellitus (DM) syndrome due to toxic misfolding of the variant proinsulin. Whereas mutations that add or remove cysteines (thereby leading to an odd number of thiol groups) generally lead to neonatal-onset DM, non-Cys-related mutations can be associated with a broad range of ages of onset. Here, we compare two mutations at a conserved position in the central B-chain α-helix: one neonatal in DM onset (Val^B18^→Gly) and the other with onset delayed until adolescence (Ala^B18^). The substitutions were introduced within a 49-residue single-chain insulin precursor optimized for folding efficiency (Zaykov, A., et al. *ACS Chem. Biol*. 9, 683-91 (2014)). Although mutations are each unfavorable, Gly^B18^ (a) more markedly perturbs DesDi folding efficiency *in vitro* than does Ala^B18^ and (b) more severely induces endoplasmic reticulum (ER) stress in cell-based studies of the respective proinsulin variants. In corresponding two-chain hormone analogs, Gly^B18^ more markedly perturbs structure, function and thermodynamic stability than does Ala^B18^. Indeed, the Gly^B18^-insulin analog forms a molten globule with attenuated α-helix content whereas the Ala^A18^ analog retains a nativelike cooperative structure with reduced free energy of unfolding (ΔΔG_u_ 1.2(±0.2) kcal/mole relative to Val^B18^ parent). We propose that mutations at B18 variably impede nascent pairing of Cys^B19^ and Cys^A20^ to an extent correlated with perturbed core packing once native disulfide pairing is achieved. Differences in age of disease onset (neonatal or adolescent) reflect relative biophysical perturbations (severe or mild) of an obligatory on-pathway protein folding intermediate.

---

Foldability is an evolved biophysical property of globular protein sequences (1). Whereas amyloid is envisaged as a polypeptide’s universal thermodynamic ground state (as a class of heteropolymers) (2,3), biological selection yields sequences capable of efficient folding to form a diversity of globular structures (4). Protection from amyloidogenesis is provided by the local thermodynamic stability of the “native state” and by kinetic barriers limiting its access to a coupled landscape of toxic aggregates (5,6). The informational content of protein sequences thus not only encodes native architectures—as visualized by the mature technologies of structural biology—but also efficient folding trajectories that can avoid kinetic traps and non-native aggregation (5,7). Residue-specific patterns of sequence conservation in a protein reflect a combination of such evolutionary constraints: some evident in the structure and function of the native state, others unseen as cryptic safeguards against off-pathway trajectories (8).

Interest in foldability and its associated safeguards has been stimulated by the recognition of diseases of protein misfolding (2,9). Classical examples are provided by extracellular amylodogenesis (10,11), pathological deposits associated with over-secretion of immunoglobulin light chains or β_2_-microglobulin (12); similar deposits comprise mutant forms of small globular domains (transthyretin and lysozyme);(13,14). Recent attention has focused on toxic misfolding within the cell (15,16). Mutations that impair folding efficiency can cause monogenic diseases even when the native state, if reached, would be capable of normal function (17). This paradigm pertains to the most common allele of cystic fibrosis (CF)^1^, which contains a deletion of a conserved Phe (Phe508) in the CFTR channel (18). A set of allele-specific small molecules (elexacaftor, tezacaftor, and ivacaftor; Trikafta [Vertex]) has been identified that restores foldability and function in such patients (a therapeutic approach given *Breakthrough* designation by the US Food and Drug Administration) (19). The key role of Phe508 in CFTR’s folding mechanism (as unmasked by medical genetics) is not apparent from structural analysis of the native state (20). Defining the cryptic roles of such conserved “safeguards” define biophysical problems of broad translational importance.

Insulin has long provided a model for the biophysical study of protein structure, stability and evolution (21-23). In the past decade a large and diverse collection of heterozygous mutations in the insulin gene (*INS*) has been found in association with non-classical diabetes mellitus (DM) (24-27). Although specific mutations can perturb essentially any step in the complex biosynthetic pathway of insulin or its hormonal function (from Golgi trafficking to prohormone processing to receptor binding)(28), the majority impair the foldability of proinsulin (the single-chain precursor of insulin (29,30)) in the rough endoplasmic reticulum (ER)(31-33). Impaired foldability of proinsulin causes ER stress (32), leading in turn to β-cell dysfunction and eventual β-cell death (34,35). Ages of disease onset of such “proinsulinopathies” from neonatal to early adulthood (31).^2^ Mutations that add or remove a Cys (thereby giving rise to an odd number of thiol groups in nascent proinsulin) ordinarily give rise to neonatal-onset DM (24,25). Dominance of the mutant allele in this syndrome reflects aberrant protein-protein interactions between the wild-type (WT) and variant proinsulins in the ER (36,37), including intermolecular disulfide pairing (38).

The present study focuses on a conserved Val in the central B-chain α-helix of insulin as a site of DM-associated clinical mutations (39,40). Val^B18^ adjoins a critical internal disufide bridge (cystine B19-A20), proposed to pair as a key early step in the folding of proinsulin (41-45). The isopropyl side chain of Val^B18^ packs within an inter-chain crevice to abut the conserved aliphatic side chains of Leu^A13^ and Leu^A16^ (Fig. 1 and Supplemental Fig. S3) (46,47). None of these five core side chains (B18, B19, A13, A16 or A20) pack at the primary hormone-receptor interface (Fig. 2) (48-50). Our interest was stimulated by the marked difference in ages of onset between variants Ala^B18^ (adolescence) (40,51)and Gly^B18^ (infancy) (39). To probe the molecular origins of this difference, we undertook a synthetic study of single-chain insulin precursors and comparative biophysical study of Val^B18^-, Ala^B18^-, and Gly^B18^-insulin analogs. Our approach was extended through cell-biological studies of ER stress on expression of the corresponding variant proinsulins.

**Figure 1.**
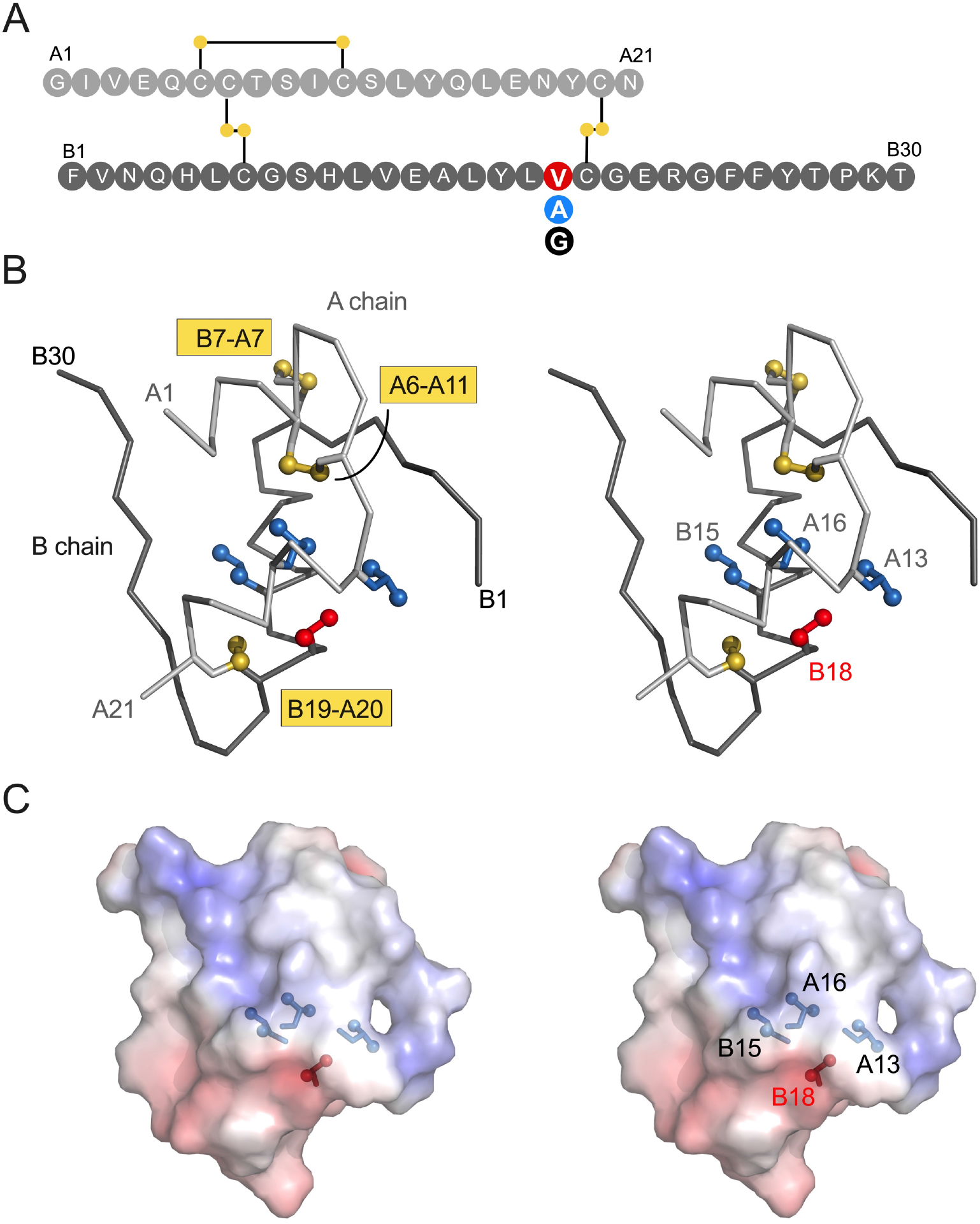
Sequence and structure of insulin. (*A*) Sequence of insulin showing clinical mutations at position B18 (highlighted by solid red circle). Residues are labeled by standard single letter code (white). The A chain is shown as light gray circles (upper sequence), and the B chain as dark gray circles (lower sequence). Neonatal onset or delayed onset is indicated by filled red or blue circles, respectively. (*B*) Stereo view of insulin monomer (C_α_-trace ribbon model; PDB entry 4INS (47)). Side chains of Leu^B15^ (blue), Val^B18^ (red), Leu^A13^ (blue), Leu^A16^ (blue) and disulfide bridges (golden) are shown as sticks. Terminal methyl groups of these side chains and sulfur atoms are shown as spheres (one-third Van der Waals radii). The A- and B chain ribbons are shown in light and dark gray, respectively. (*C*) Spatial environments of residue B18 (color coding and drawing same as in panel B) shown as electrostatic surfaces that are calculated in the absence of B18 side chain. blue and red surfaces are coded by positive or negative electrostatic potential.

**Figure 2.**
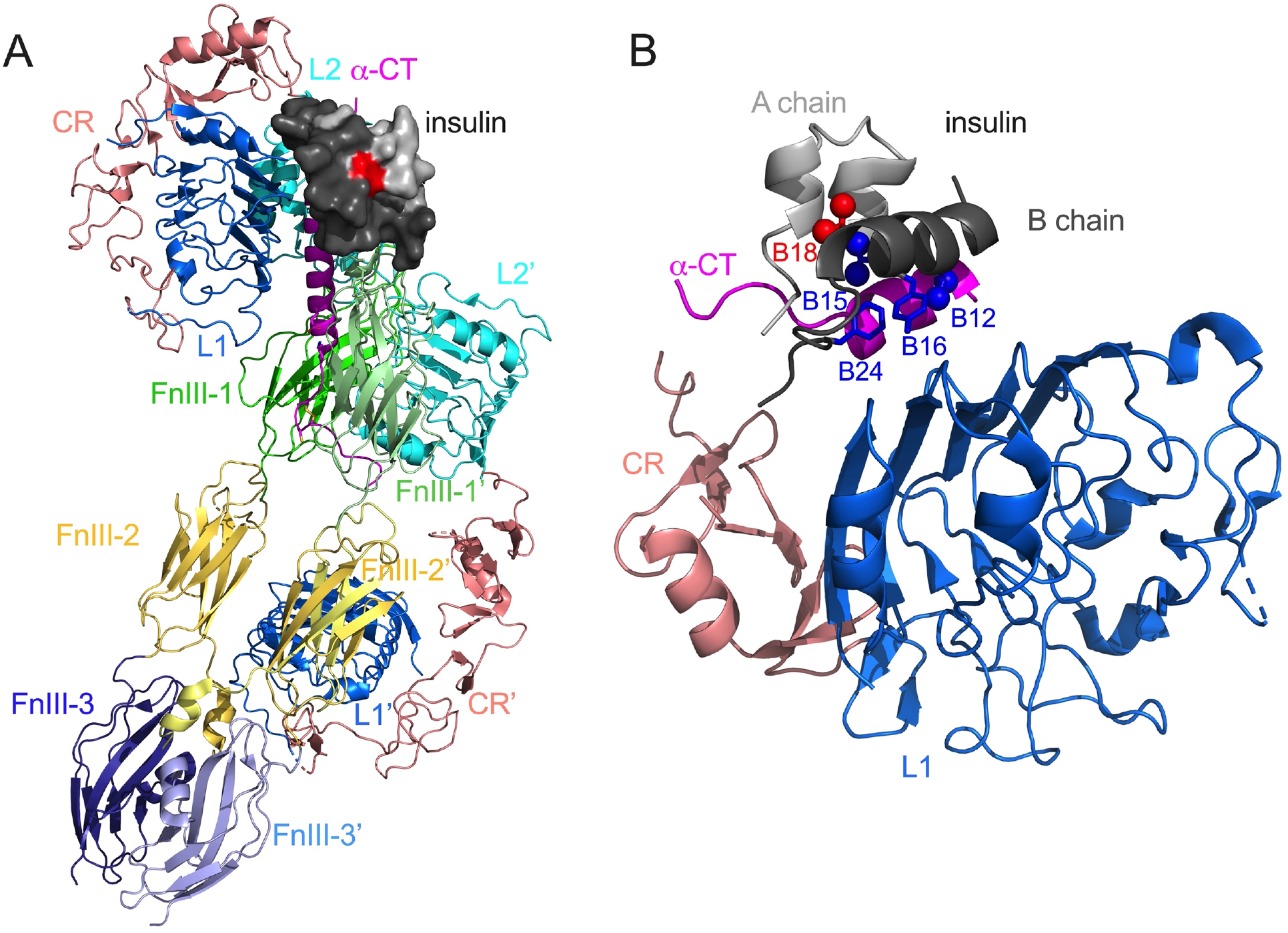
Cryo-EM structures of insulin receptor. (*A*) Structure of receptor ectodomain (PDB entry 6HN5 for upper domain and 6HN4 for lower domain (50)). Insulin is shown as surface (A chain, light gray; B chain, dark gray). Val^B18^ highlighted in red. (*B*) Structure of microreceptor (PDB enry 4OGA (49)). Receptor bound insulin shown as cartoon (color coding same as in panel *A*). Residues Phe^B24^, Tyr^B16^, Val^B12^ and Leu^B15^ are shown in blue; Val^B18^ shown in red. Methyl groups of these side chains are shown as spheres. In each case the domains of receptor are presented as cartoons and are labeled within the figure.

To our knowledge, this study represents the first interrogation of a cryptic inter-chain crevice in the insulin molecule. Our findings suggest that native packing of branched aliphatic side chains in this potential space is critical to the foldability of proinsulin, presumably by promoting nascent pairing of cystine B19-A20. These conserved side chains also buttress and stabilizes the mature hormone’s receptor-binding surface. Age of DM onset and ER stress thus correlate with both extent of impaired foldability *in vitro* and extent of structural perturbation in the native state, once reached. We envisage that this correlation will apply generally in the mutant proinsulin syndrome (52).

## Results

### Chemical synthesis provides an assay of foldability

To investigate the folding efficiencies of variant insulin analogs, we have chosen DesDi as a synthetic model (53). DesDi (a mini-proinsulin) is a single-chain 49-residue polypeptide t with substitution Pro^B28^→Lys^3^ and a Lys^B28^-Gly^A1^ peptide bond; it hence lacks residues B29-B30. Lys^B28^ provides a cleavage site (for a Lys-specific endoprotease) to yield an active two-chain insulin analog (Lys^B28^-*des*-dipeptide[B29,B30]-human insulin). DesDi exhibits higher folding efficiency than proinsulin or other mini-proinsulins, presumably due to optimization of the B28-A1 linkage to bias the unfolded-state ensemble toward on-pathway conformations. This template thus enables preparation of mutant insulin analogs not otherwise amenable to insulin chain combination (22,44).

Peptides were obtained by solid-phase peptide synthesis (SPPS) with a heat-assisted coupling/deprotection protocol (see Experimental Procedures and Supplemental Methods). The crude polypeptides were subjected to oxidative folding reactions to assess relative yields. The parent DesDi (with Val^B18^) thus provided a baseline for comparative studies of the clinical variants (Ala^B18^ and Gly^B18^). Folding was monitored by analytical reverse-phase high-performance liquid chromatography (rp-HPLC). Ala^B18^ DesDi exhibited a ∼twofold reduction in folding efficiency whereaas Gly^B18^ exhibited ∼eightfold reduction (Fig. 3). The ratio of yields following semi-preparative rp-HPLC purification followed the same trend (See Supplemental Table S1).

**Figure 3.**
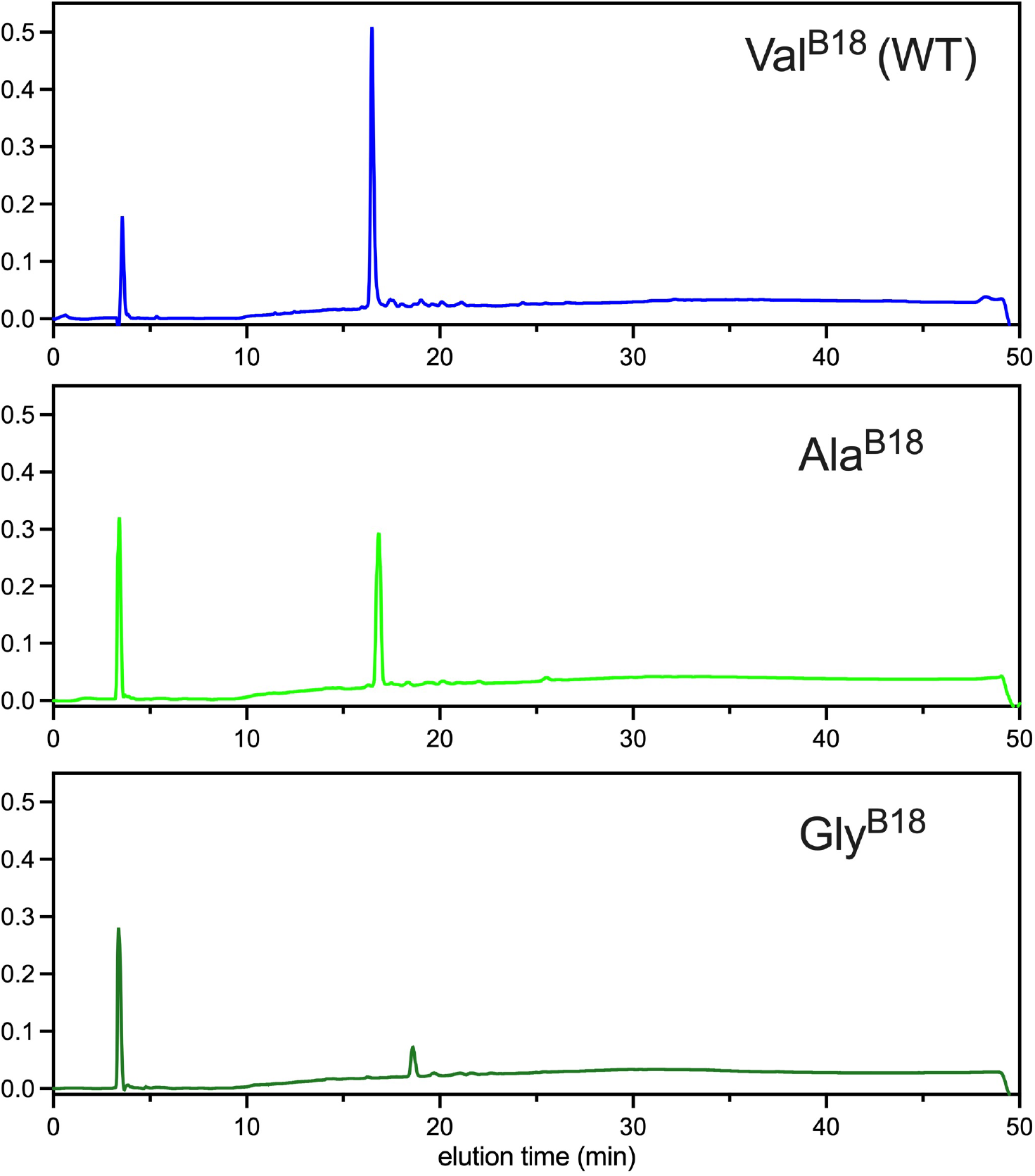
Comparison of folding profiles of wild-type (Val) and clinical mutations (Ala and Gly) at position B18 using DesDi synthetic precursor. Folding reactions were carried out using crude polypeptides at 100 mg scale (∼18 δmol) at 0.1 mM peptide concentration in 20 mM glycine, 2 mM Cysteine at pH 10.5 aqueous buffer under atmosphere of air (4 °C).

Once folded, the three single-chain precursors were treated with Endo Lys-C endoprotease to obtain corresponding two-chain *des*-dipeptide[B29, B30] insulin analogs (Supplemental Fig. S4-S9). Although Lys^B28^-*des*-dipeptide[B29,B30]-insulin is less stable than WT insulin (and impaired in dimerization), this template enabled comparison of B18 side chains (Val, Ala or Gly) with respect to relative changes in structure, stability and function.

To gain additional insight into *in-vitro* foldabilities (44,54), we next exploited chain combination. The pairing reaction of isolated A- and B chains is ordinarily tolerates a wide variety of substitutions: over the past five decades hundreds of insulin analogs have been so prepared, demonstrating that the information required for proinsulin folding is contained within the A- and B-chain sequences (55). Under the present reaction conditions (see Experimental Procedures and Supplemental Methods), the WT chains (as S-sulfonate derivatives) combined to yield WT insulin in expected yield. In striking contrast, combination of the WT A chain with either of the two variant B chains (Ala^B18^ and Gly^B18^) each gave only trace product (Supplemental Fig. S13-15). Although this failure of chain combination highlights the present mutational perturbations, relative yields did not correlate with respective ages of onset, presumably due to the lower baseline robustness of chain combination relative to the folding of single-chain precursors.

### CD studies uncover mutational perturbations

Far-ultraviolet (UV) circular dichroism (CD) spectra of single-chain analogs essentially had a similar pattern in 200-255 nm range except for Gly^B18^ which had attenuated 195 nm peak (Fig. 4*A*). Deconvolution by SELCON algorithm (56) indicated these analogs with similar helical propensities and Ala being slightly better compared to Val or Gly. (See Supplemental Table S2) Free energies of unfolding at 37 °C as inferred from two-state modeling of chemical denaturation indicated the decremental values in the order Val (more stable) > Ala > Gly (less stable) (Fig. 4*B*). Noticeable changes were seen in spectra of respective two chain analogs (Fig. 4*C*). While Gly^B18^ found to be completely disordered, Ala^B18^ had better helical structure even compared to Val^B18^ presumably due to its superior helical propensity when compared to native valine (57,58) (See Table 2 and Supplemental Table S2). SELCON results clearly showed the Gly^B18^ being unstructured with 20.6% of α-helical content compared to Val^B18^ (45.2%) or Ala^B18^ (47.5%). (Table S2). Guanidine denaturation studies of two chain analogs at 25 °C indicated decreased thermodynamic stabilities for Ala^B18^ analog compared to Val^B18^ (ΔG_u_ 1.2 (±0.2) kcal/mole] (Table 1). Gly^B18^ had a parabolic transition indicative of molten globule like structure (Fig. 4*D*).

**Figure 4.**
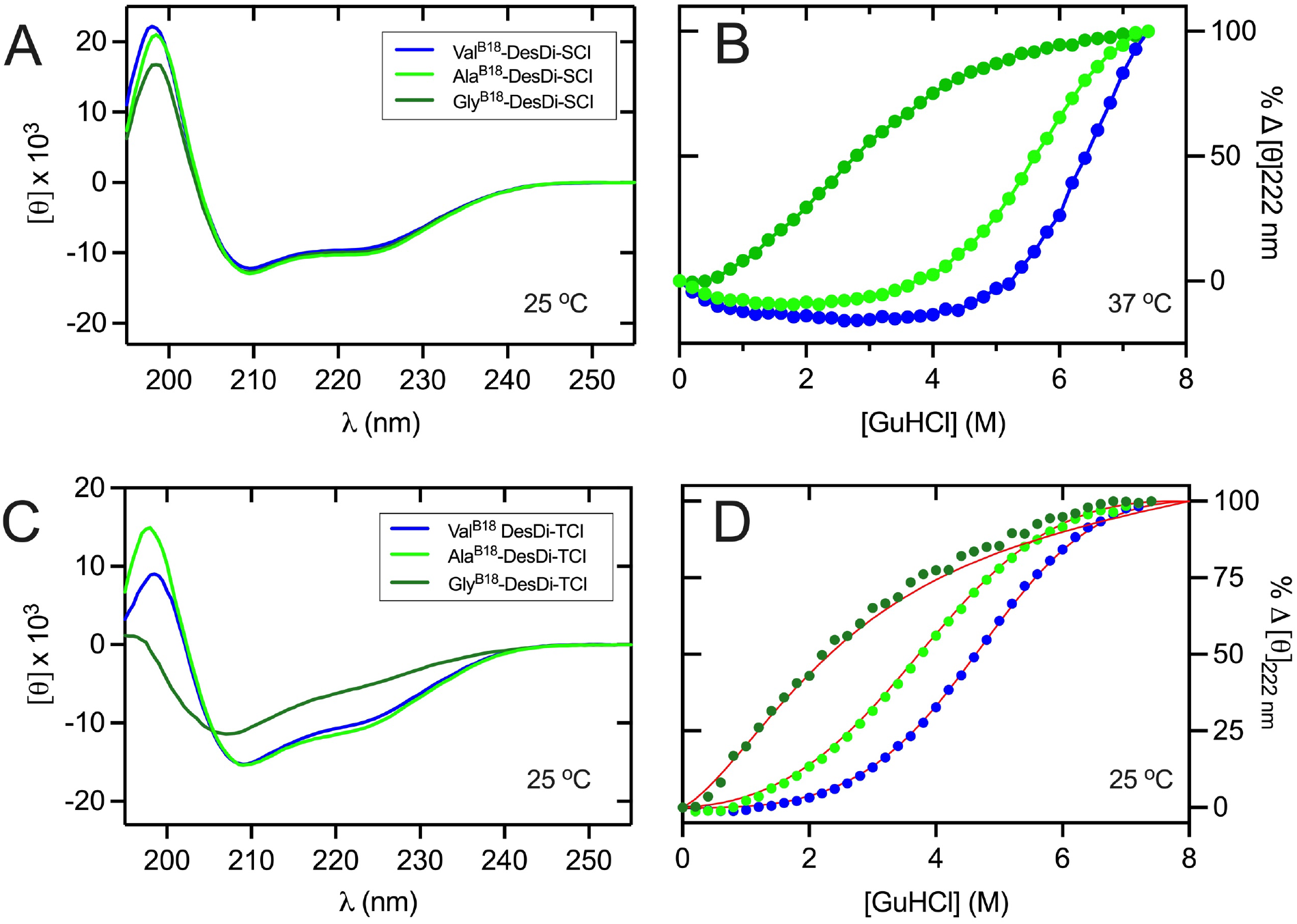
CD and guanidine denaturation studies of DesDi insulin analogs. (*A, B*) Far-UV spectra of Val^B18^ (WT), Ala^B18^, and Gly^B18^-DesDi single-chain insulin analogs (*A*) and corresponding CD-detected unfolding transitions monitored by ellipticity at 222 nm (*B*). (*C, D*) Spectra of respective two chain analogs (*C*) and corresponding unfolding transitions (*D*). Color codes are defined in panels A and C. All the studies were performed at 25°C except for the guanidine denaturation studies of single-chain precursors which was done at 37 °C owing to their relative higher stability at 25 °C. The resulting stability values are given in Table 1.

**Table 1.** CD denaturation studies

| *analog* | $\Delta G_u$ (kcal/mol)<sup>a</sup> | *m*-value<sup>b</sup> | $C_{mid}$ (M)<sup>c</sup> |
| --- | --- | --- | --- |
| *single-chain precursors (37 °C)* |  |  |  |
| DesDi-Val<sup>B18</sup> (parent) | 7.5 ± 0.4<sup>d</sup> | 1.10 ± 0.06 | 6.8 ± 0.4 |
| DesDi-Ala<sup>B18</sup> | 4.0 ± 0.1 | 0.60 ± 0.03 | 6.7 ± 0.2 |
| DesDi-Gly<sup>B18</sup> | 0.6 ± 0.1 | 0.49 ± 0.02 | 1.2 ± 0.1 |
| *two-chain insulin analogs (25 °C)* |  |  |  |
| human insulin | 3.3 ± 0.1 | 0.69 ± 0.01 | 4.8 ± 0.1 |
| insulin *lispro* | 3.0 ± 0.1 | 0.63 ± 0.01 | 4.7 ± 0.1 |
| DesDi-Val<sup>B18</sup> (WT) | 2.9 ± 0.1 | 0.64 ± 0.02 | 4.5 ± 0.1 |
| DesDi-Ala<sup>B18</sup> | 1.7 ± 0.1 | 0.46 ± 0.01 | 3.7 ± 0.1 |
| DesDi-Gly<sup>B18</sup> | N.D.<sup>e</sup> | N.D. | N.D. |

<sup>a</sup>Estimates of free-energy changes were inferred from two-state modeling of chemical-denaturation titrations as described (23); assays were performed in 10 mM potassium phosphate (pH 7.4) and 50 mM KCl.
<sup>b</sup>The *m*-value is the slope of unfolding free energy $\Delta G_u$ versus molar concentration of denaturant
<sup>c</sup> $C_{mid}$ is the guanidine denaturant concentration at which 50% of the protein is unfolded.
<sup>d</sup>The parent single-chain analog (Lys<sup>B28</sup>-*des*-dipeptide[B29,B30]-miniproinsulin (53)) is too stable to unfold completely in 8 M guanidine-HCl and 37 °C; this limited modeling of the post-transition baseline and introduced a larger uncertainty in model parameters.
<sup>e</sup>This insulin analog exhibited a non-cooperative unfolding transition.

**Table 2.**
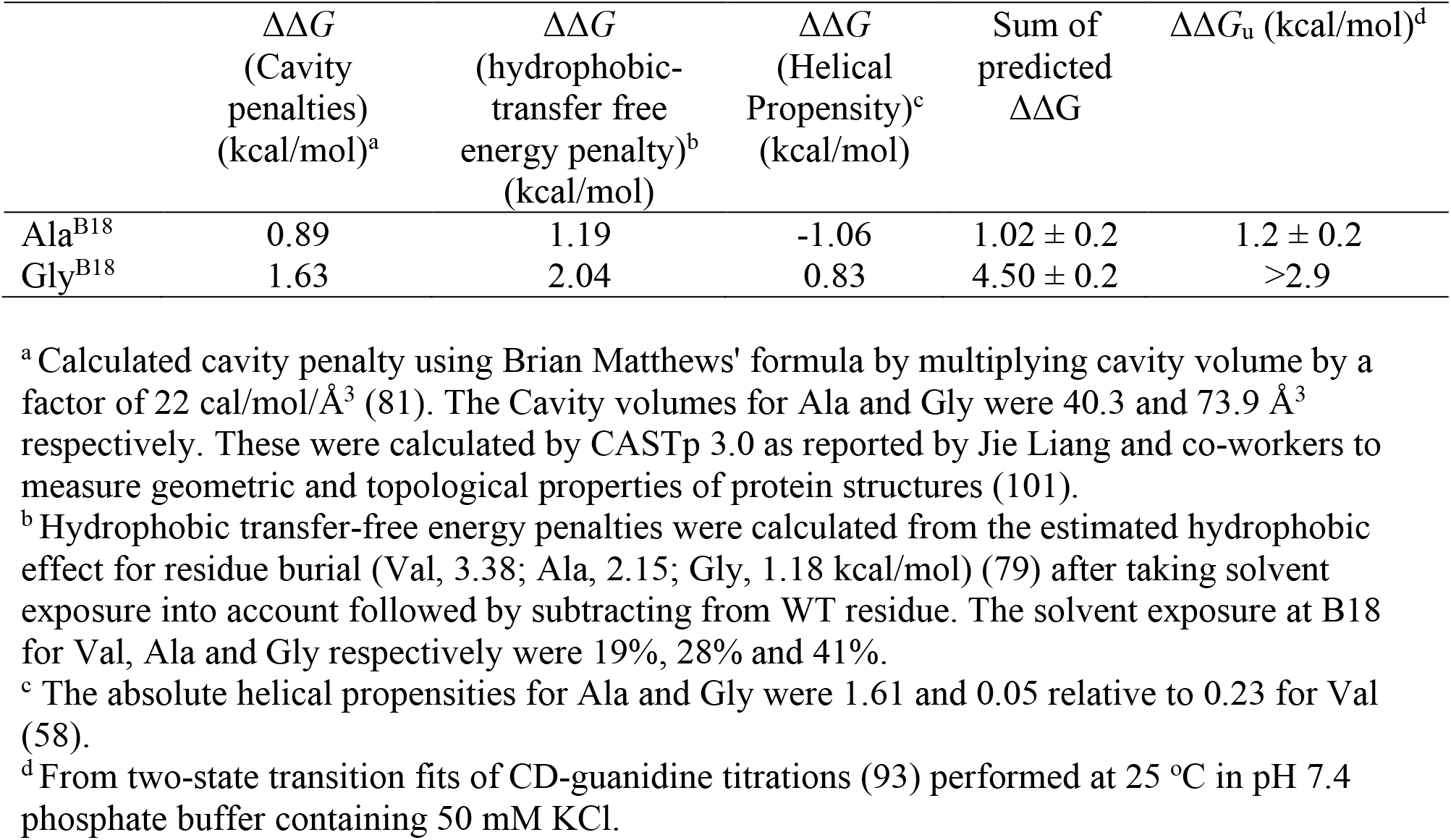
Comparison of predicted and observed thermodynamic parameters.

| | $\Delta\Delta G$<br>(Cavity<br>penalties)<br>(kcal/mol)<sup>a</sup> | $\Delta\Delta G$<br>(hydrophobic-<br>transfer free<br>energy penalty)<sup>b</sup><br>(kcal/mol) | $\Delta\Delta G$<br>(Helical<br>Propensity)<sup>c</sup><br>(kcal/mol) | Sum of<br>predicted<br>$\Delta\Delta G$ | $\Delta\Delta G_u$ (kcal/mol)<sup>d</sup> |
| --- | --- | --- | --- | --- | --- |
| Ala<sup>B18</sup> | 0.89 | 1.19 | -1.06 | 1.02 ± 0.2 | 1.2 ± 0.2 |
| Gly<sup>B18</sup> | 1.63 | 2.04 | 0.83 | 4.50 ± 0.2 | >2.9 |

<sup>a</sup> Calculated cavity penalty using Brian Matthews' formula by multiplying cavity volume by a factor of 22 cal/mol/Å<sup>3</sup> (81). The Cavity volumes for Ala and Gly were 40.3 and 73.9 Å<sup>3</sup> respectively. These were calculated by CASTp 3.0 as reported by Jie Liang and co-workers to measure geometric and topological properties of protein structures (101).
<sup>b</sup> Hydrophobic transfer-free energy penalties were calculated from the estimated hydrophobic effect for residue burial (Val, 3.38; Ala, 2.15; Gly, 1.18 kcal/mol) (79) after taking solvent exposure into account followed by subtracting from WT residue. The solvent exposure at B18 for Val, Ala and Gly respectively were 19%, 28% and 41%.
<sup>c</sup> The absolute helical propensities for Ala and Gly were 1.61 and 0.05 relative to 0.23 for Val (58).
<sup>d</sup> From two-state transition fits of CD-guanidine titrations (93) performed at 25 °C in pH 7.4 phosphate buffer containing 50 mM KCl.

Temperature-dependent CD spectra indicated no significant changes in the 200-255 nm range for single-chain analogs (See supplemental Fig. S16, left panel). The peak at 195 nm however found to attenuate going from 4 °C to 25 °C to 37 °C. In the case of two chain analogs (supplemental Fig. S16, right panel), noticeable changes were seen in Gly^B18^ where disordered structure at 25 °C or 37 °C gained significant structural features at 4 °C. Whereas Val^B18^ and Ala^B18^ showed marginal effect with temperature.

### NMR spectra probe hydrophobic-core residues

Two chain versions of DesDi-Val^B18^, Ala^B18^ and Gly^B18^ were analyzed by 1D ^1^H-NMR spectra, 2D TOCSY, NOESY and ^1^H-^13^C HSQC correlation spectra to gain structural insights. The assignments of ^1^H and ^13^C chemical shift were established through unique through-bond connections in TOCSY and specific NOE pattern in NOESY spectra, and confirmed in ^13^C-HSQC correlation spectra (Fig. 5 and 6; also see Supplemental Fig. S17). The Val^B18^ analog exhibits ^1^H-NMR spectral property of native-like insulin as observed in the aromatic region and in the upfield-shifted methyl region (Fig. 5, *far right*). The well-resolved methyl resonances indicated as a: Ile^A2^ γ_2_-CH_3_; b: Ile^A10^ γ_2_-CH_3_; c: Leu^B15^ δ_2_-CH_3_ d: Ile^A2^ δ_1_-CH_3_; e: Ile^A10^ δ_1_-CH_3_; f: Leu^B15^ δ_1_-CH_3_. The signature long-range NOEs from the aromatic protons of Phe^B24^ and Tyr^B26^ residues to methyl protons of Leu^B15^ residue, and that from the aromatic protons of Tyr^A19^ to methyl protons of Ile^A2^ and Leu^A16^, were also observed, indicating native-like conformation of DesDi-Val^B18^ (Fig. 6*A,D*). In comparison to DesDi-Val^B18^, 1D and 2D homonuclear ^1^H-NMR spectra of DesDi-Ala^B18^ indicated similar, but not identical, chemical-shift dispersion and typical native-like packing of hydrophobic residues in the hydrophobic core of insulin (Fig. 5, *middle panel*). The secondary ^1^H-NMR chemical shifts of the upfield-shifted side chain of Leu^B15^ (a characteristic signature of the Phe^B24^ ring current in the native B-chain super-secondary structure) and that of Phe^B24^ aromatic protons were attenuated (shifted toward random-coil direction) in the Ala^B18^ variant. Similar signature long-range NOEs between methyl protons and those of the aromatic side chain were also observed, except signal broadening related to Phe^B24^ probably due to presumptive conformational exchange or self-association (Fig. 6*B,E*). 2D heteronuclear ^1^H-^13^C HSQC correlation spectrum can provide structural footprint at the atomic level. Patterns of nature abundance ^1^H-^13^C HSQC NMR spectra of DesDi-Val^B18^ and DesDi-Ala^B18^ also exhibit typical spectral characteristics of native-like insulin both in aromatic and methyl region (See Supplemental Fig. S17), as observed in 1D and 2D ^1^H-NMR spectra. These qualitative ^1^H and ^13^C spectra features provide evidence of a native-like structure of DesDi-Ala^B18^ with a local change in structure and/or dynamics near the site of substitution transmitted to the central B-chain α-helix and to β-strand portion of the B-chain. The DesDi-Gly^B18^ lost its secondary chemical shift dispersion in both aromatic and methyl region (Fig. 5, *top panel*) and the signature long-range NOEs between methyl and aromatic protons were not observed (Fig. 6*C,F*), suggesting a unfolded structure.

**Figure 5.**
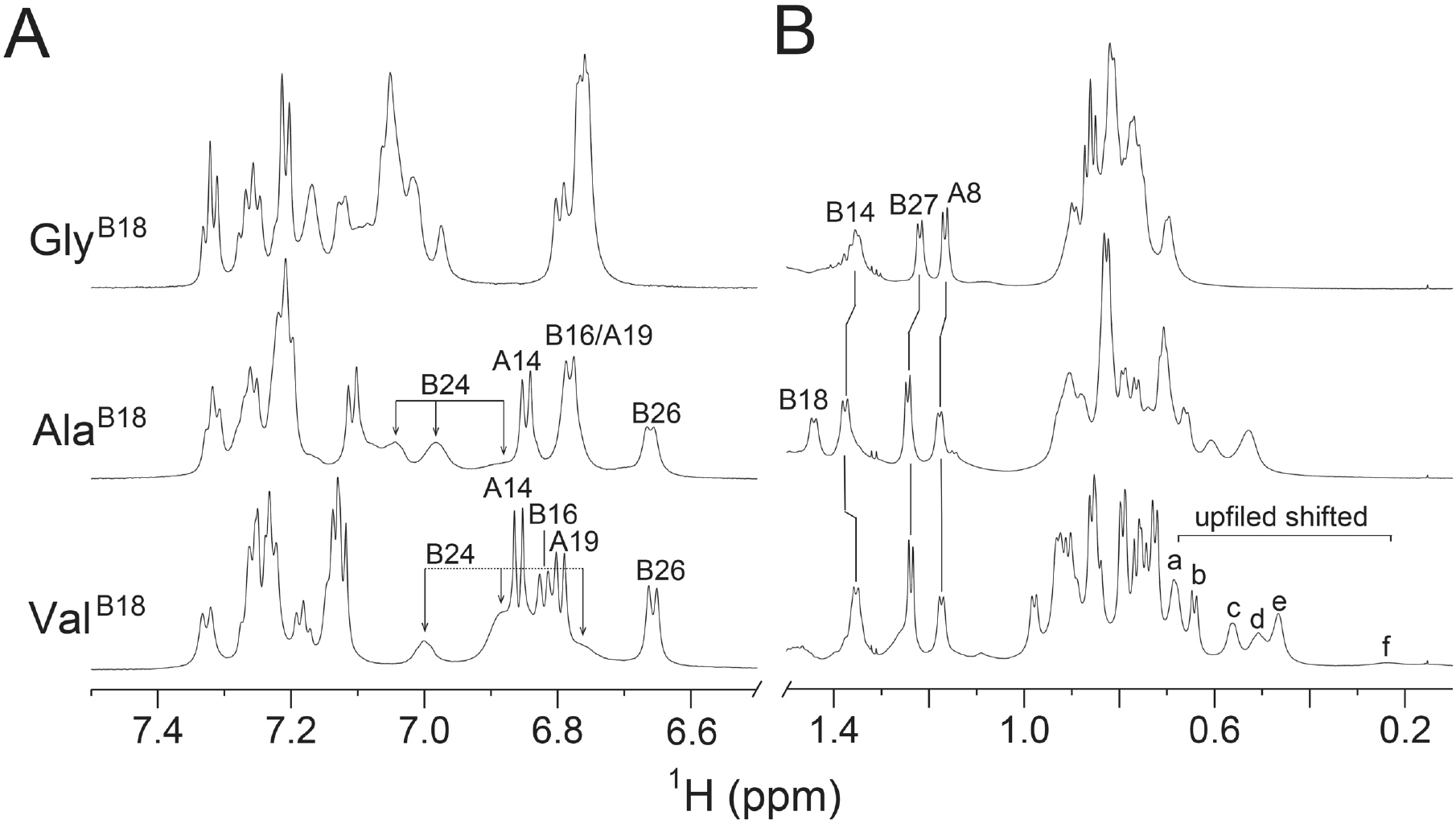
Stack plot of 1D ^1^H-NMR spectra of DesDi insulin analogs. (*A*) aromatic and (*B*) methyl region. Bottom panel is parent DesDi Val^B18^ analog, middle panel is Ala^B18^ analog and top panel is Gly^B18^ analog. The Val^B18^ analog exhibits spectral property of native-like insulin as observed in the aromatic region and in the upfield-shifted methyl region (*far right*). The well-resolved methyl resonances indicated as a: Ile^A2^ γ_2_-CH_3_; b: Ile^A10^ γ_2_-CH_3_; c: Leu^B15^ δ_2_-CH_3_ d: Ile^A2^ δ_1_-CH_3_; e: Ile^A10^ δ_1_-CH_3_; f: Leu^B15^ δ_1_-CH_3_. For Ala^B18^ analog, aromatic and methyl resonances of signature residues Phe^B24^ and Leu^B15^ shifted to downfield, as well as exhibited a reduction in chemical-shift dispersion. Spectra were acquired at pD 7.4 (direct meter reading) at 32 °C in D_2_O.

**Figure 6.**
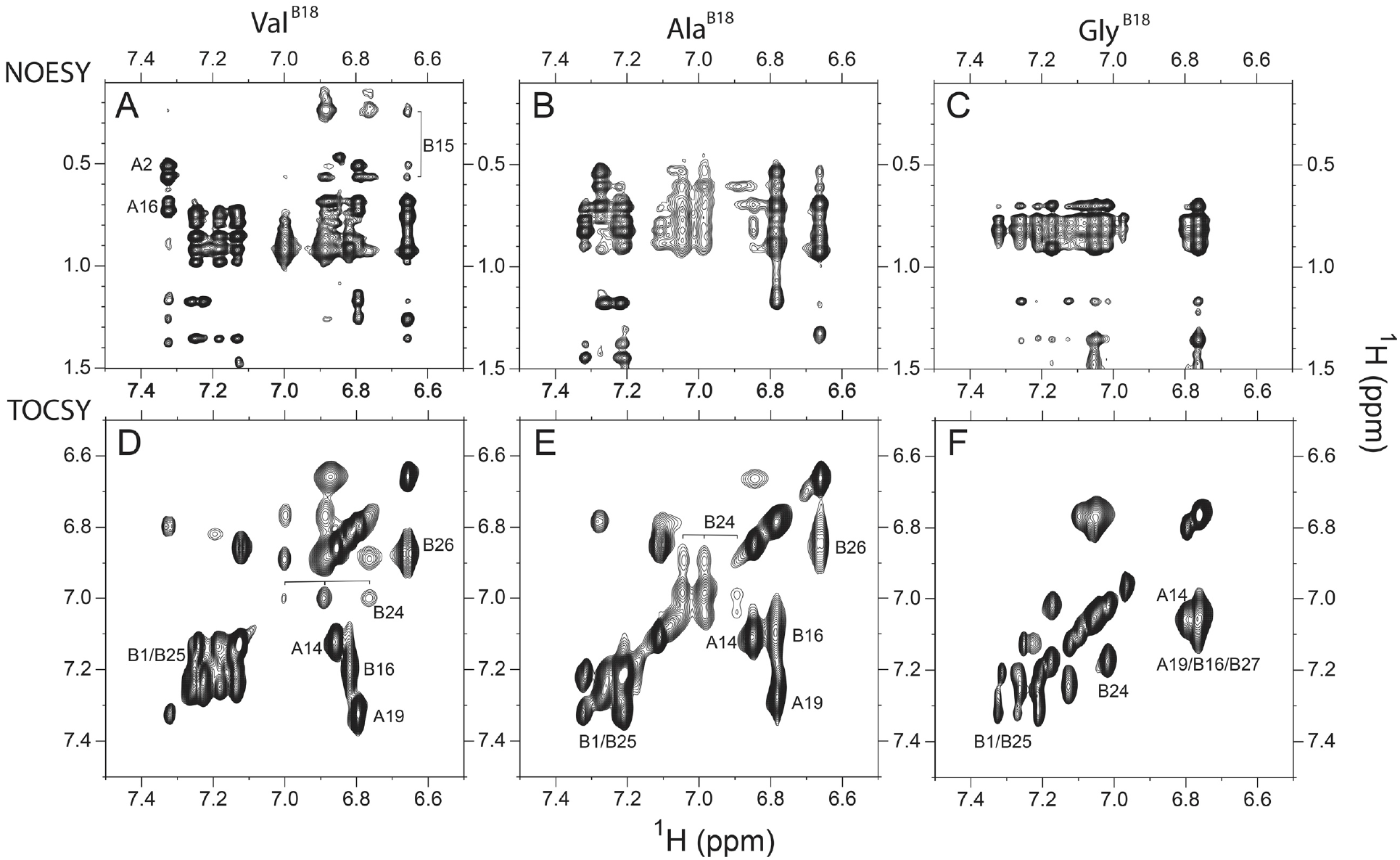
2D NMR studies. Top panel shows NOEs from aromatic to aliphatic protons in 2D NOESY spectra of DesDi Val^B18^ analog (*A*), Ala^B18^ analog (*B*) and Gly^B18^ analog (*C*). Bottom panel shows aromatic resonances of 2D TOCSY spectra of DesDi Val^B18^ analog (*D*), Ala^B18^ analog (*E*) and Gly^B18^ analog (*F*). Spectra were acquired at pD 7.4 (direct meter reading) at 32 °C in D_2_O.

The temperature-dependent NMR spectra were also acquired at 12, 22 and 32 °C, respectively (See Supplemental Fig. S18). The chemical shift dispersion of DesDi-Ala^B18^ and DesDi-Gly^B18^ analogs slightly increased when lowering the temperature. Interestingly, one most upfield shifted peak in the DesDi-Ala^B18 1^H-NMR spectrum was observed at 12 °C around 0.27 ppm, possibly from Leu^B15^ δ_1_-CH_3_ and/or Ile^A10^ γ_12_-CH_2_ protons. Due to the spectral overlap and line broadening of Leu^B15^-related resonances at 12 °C, it was difficult to assign. Therefore, the ^1^H-NMR spectrum of DesDi-Ala^B18^ was acquired at ∼9-times lower concentration at 32 °C (See Supplemental Fig. S19). Well-resolved and upfield-shifted signals were observed in the concentration of ∼0.03 mM and the spectral pattern both in aromatic and methyl region was very similar to that of DesDi-Val^B18^ analog (Supplemental Fig. S19). Those peaks shifted toward upfield and the chemical shift dispersion increased upon temperature decrease.

### Age of disease onset correlates with ER stress

The endoplasmic reticulum (ER) is the central site for protein folding, post-translational modifications, and transportation (59). The accumulation of misfolded proteins in the ER would induce the expression of glucose-regulated proteins (GRPs) including GRP78/heavy chain binding protein (BiP) and enhanced the expression of the C/EBP Homologous Protein (CHOP) (60). This response is regulated by the signaling pathway initiated by the phosphorylation of PERK (Fig. 7*A*).

**Figure 7.**
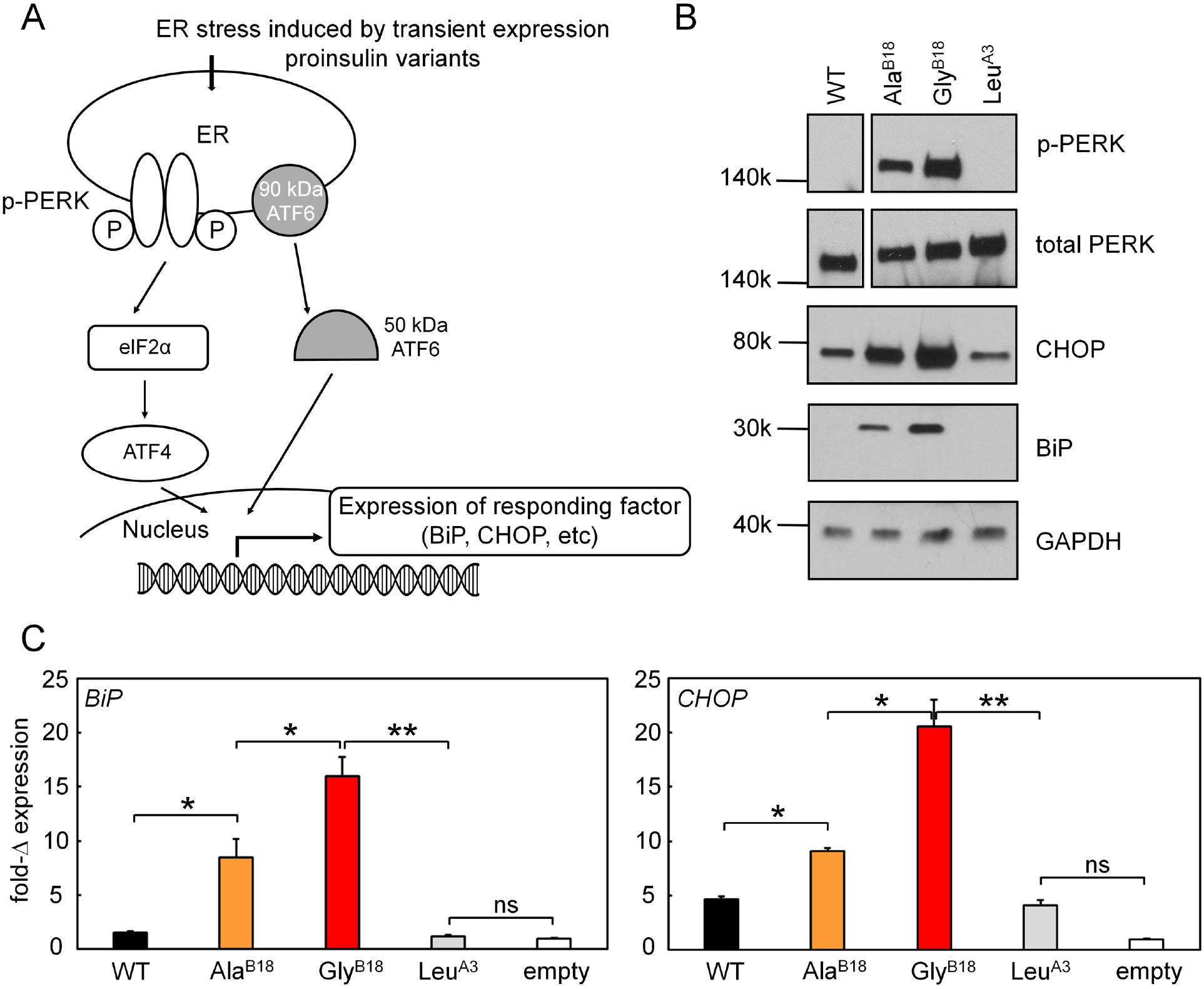
Proinsulin mutants induce the ER stress. (*A*) Schematic figure presenting the ER stress induced by expression of proinsulin analogs in cells (102). Mutant proinsulin variants give rise to accumulation of unfolded protein in the ER resulting in stress, which causes phosphorylation of PERK and induction of ISR. CHOP induction is a downstream response of the ISR and BiP is a chaperon, which would be activated and lead to its increased expression. The protein levels of p-PERK and the accumulations of ER stress markers BiP and CHOP were tested by Western blot approach (*B*). The transcription responses of these two ER stress markers were also monitored by qPCR assay (*C*). (*B*) Western-blot assays probing the ER stress markers: p-PERK/PERK, CHOP, and BiP, induced by the expression of proinsulin variants. Expression of WT and Leu^A3^ proinsulin provides the controls showing inactive ER stress. (*C*) Real-time qPCR assay probing the transcription responses of *BiP* and *CHOP* genes induced by WT and variants of proinsulin. Gene markers for ER stress were significantly activated by the expressions of Gly^B18^ (red) and Ala^B18^ (orange) proinsulin. WT and Leu^A3^ proinsulin served as controls. Asterisks (*) and (**) indicate p-value < 0.05 and < 0.01. The “ns” indicates p-value > 0.05.

To examine the ER stress effect induced by misfolded proinsulin-driven signaling pathway in mammalian cells, the HEK 293T cell lines were transiently transfected with WT (Val^B18^) and two of mutations (Ala^B18^ and Gly^B18^), we investigated the accumulation of ER stress markers in protein level. The phosphorylated PERK kinase exhibited significantly increasing in cells expressing Ala^B18^ and Gly^B18^ proinsulin. The accumulations of the other two representative marker proteins, CHOP and BiP (61), also increased comparing to the WT proinsulin control and a function-defected mutation without structural impairment (62,63) (Leu^A3^; Fig. 7*B*). The ER stress also activates *BiP* promoter leading to its increased mRNA transcription to resolve the stress by properly folding these proinsulins (61). Since these mutated insulin proteins still fail to be folded properly, the Integrated Stress Response (ISR) is induced to attenuate global protein synthesis and the *CHOP* gene is activated for triggering apoptosis. Our result shows that the gene expressions of CHOP and BiP exhibit significantly increasing (Fig. 7*C*), which is consistent with the phenomenon of ER stress responses.

### Mutations at B18 perturb hormonal activity

Insulin’s hinge-opening at a dimer-related interface is coupled to closure of the insulin receptor (IR) ectodomain legs. This engagement leads to TK trans-phosphorylation and receptor activation. Activation of signal pathway was probed by a kinase cascade in HepG2 cells. Quantitative dose-dependent IR autophosphorylation was evaluated in a 96-well plate assay (Fig. 8*A,B*). Insulin *lispro* (KP)^[2]^ and Val^B18^ DesDi (WT) insulin triggered a robust autophosphorylation readouts whereas the mutations of Ala^B18^ and Gly^B18^ DesDi exhibited significantly reduced activities. These data indicate that the requirement of high-dose exposure of Ala^B18^ and Gly^B18^ DesDi analogs to reach native-like activities (>3000 nM, comparing to KP and WT DesDi for only 50 nM of treatment; Fig. 8*B*). An overview of phosphorylation of IR and downstream phosphorylation of Ser-Thr protein kinase AKT (protein kinase B; ratio p-AKT/AKT) at different hormone doses (50 nM and 3000 nM) were provided by western blot (Fig. 8*C*). These data support the findings that these two mutations retain little activities and at least 60-fold higher doses are required for comparable IR-activation function.

**Figure 8.**
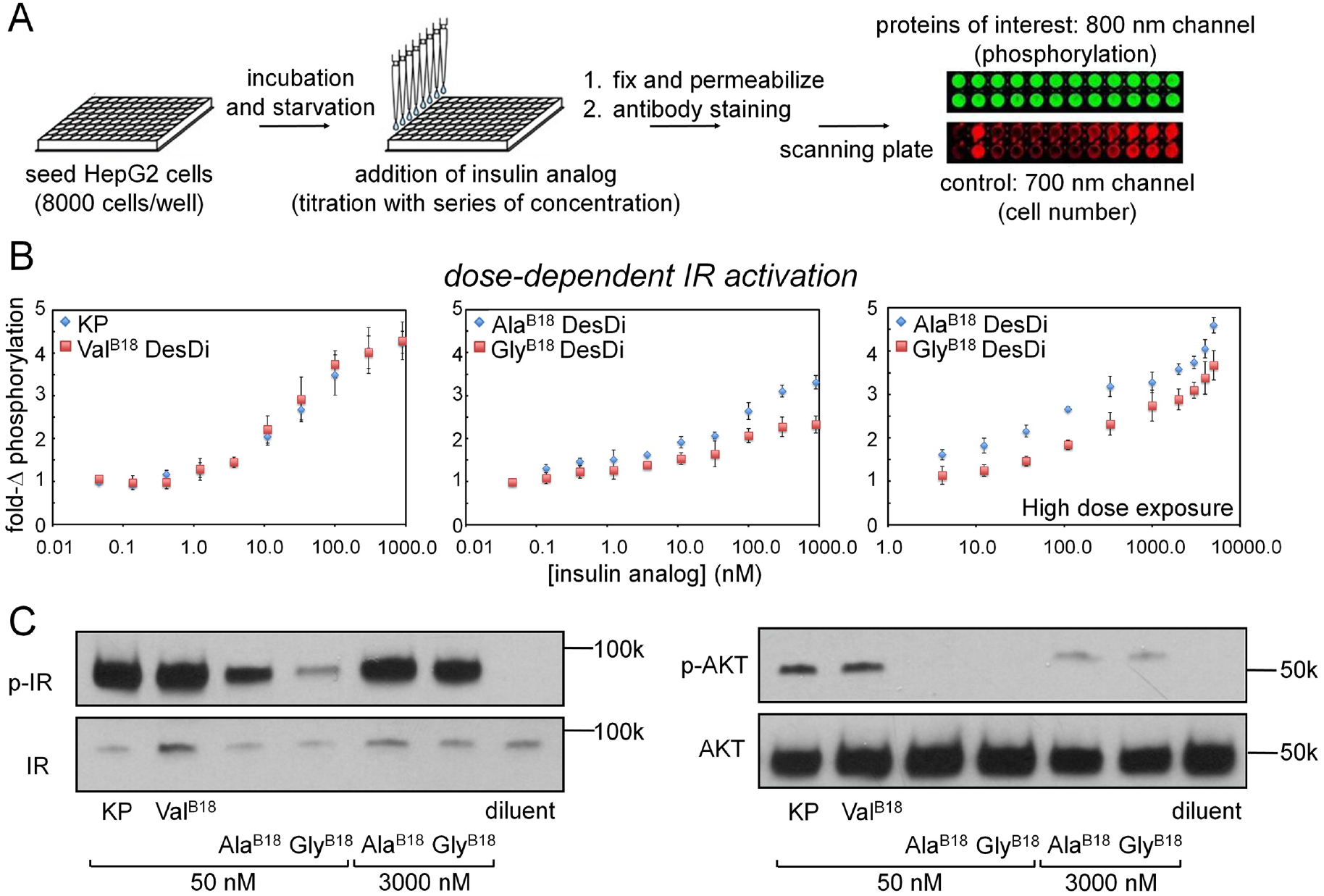
Dose-response studies of insulin activity. (*A*) Schematic outline of in-cell assays probing IR autophosphorylation in exposure of various insulin analogs. Flow chart shows the procedure to monitor hormone-induced IR signaling via in-cell illumination assay using the ratio of readouts between 800 nm (phosphorylation) and 700 nm (cell number control). (*B*) The IR autophosphorylation on hormone binding. Insulin lispro (labeled as KP) and Val^B18^ (WT) DesDi insulin share similar activity in all the titration conditions (left). Mutations (Ala and Gly) in B18 position significantly reduced the activities (middle) and only reach the WT DesDi-like activities when exposed in high dose condition (>3000 nM; right). (*C*) Western-blot assays probing the p-IR/IR and p-AKT/AKT in the ordinary (50 nM) and high dose (3000 nM) of analog Ala^B18^ and Gly^B18^ in medium.

The *in vivo* potencies of these analogs, evaluated by subcutaneous (SQ) injection in diabetic rats (Fig. 9) indicated Ala^B18^ as having reduced blood glucose lowering capability when compared to Val^B18^ (p-value <0.05) (Fig. 9*A,B*). On the other hand, Gly^B18^ had very low activity even at twofold or fourfold higher doses per kilogram (Fig. 9*C,D*). Although the individual p-values were 0.07 and 0.22, respectively, combining the data supported that hypothesis that Gly^B18^ impairs activity with p < 0.05. It is not known whether the patients with these monogenic diabetes syndromes secrete the mutant insulin (39,40) and thus whether their reduced potencies contribute to disease onset of progression.

**Figure 9.**
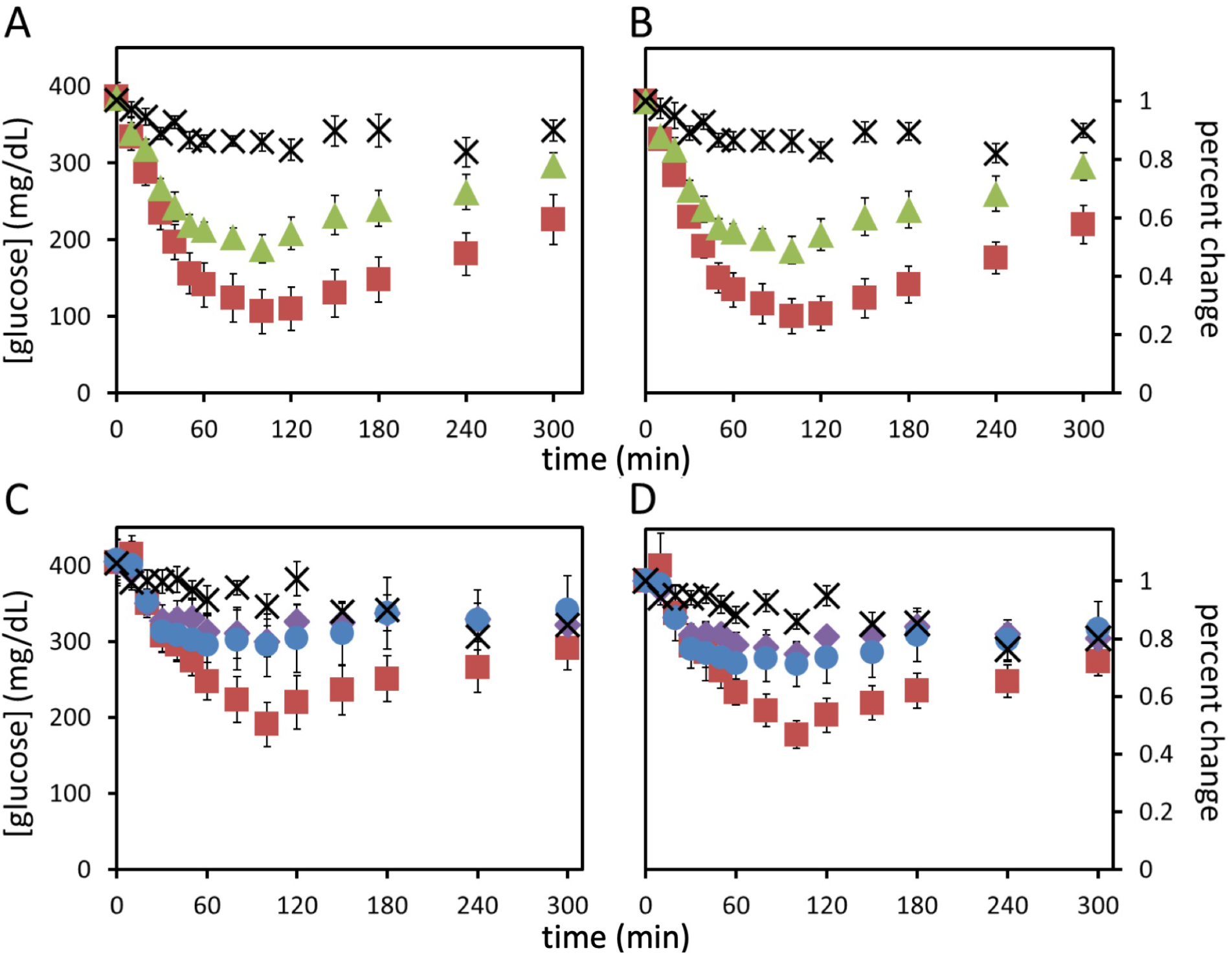
Rat studies of DesDi two chain insulin analogues. (*A*) time course of [blood glucose] following SQ injection. Val^B18^ (red squares, n=6); Ala^B18^ (green triangle, n=6); diluent (X, 100 μL, n=6). The dose of WT or mutant insulin was 6 nmol/kg rat. (*C*) Dose-response studies of Gly^B18^ relative to parent Val^B18^ analog at a 6 nmol/kg rat dose (red squares; N=5). Doses of the Gly^B18^ analog: 12 nmol/kg rat (blue circles; N=5) and 24 nmol/kg rat (purple diamond; N=5). (*B*) and (*D*), normalized blood glucose profiles for the data in panels *A* and *C*, respectively. Standard errors of mean are given by black vertical error bars. Quantitative and statistical analyses are provided in Supplemental Fig. S14.

## DISCUSSION

Insulin sequences have been highly conserved in the evolutionary radiation of vertebrates: critical roles in fetal development and metabolic homeostasis have evidently imposed rigorous functional selection for more than 500 million years (64,65). Such conservation is particularly marked in the hydrophobic core wherein two internal disulfide bridges (cystines B19-A20 and A6-A11) nestle among the invariant side chains of Leu^B6^, Leu^B15^, Phe^B25^, Ile^A2^, Leu^A16^ and Tyr^A19^ (47). This invariant framework is shared by insulin-like growth factors (IGF-1 and IGF-2), homologs of proinsulin arising from gene duplication in the transition from protochordates at an early stage of vertebrate evolution (65,66). Purifying selection in this family is envisioned at general steps in protein biosynthesis and function: from oxidative folding in the ER (enabling specific disulfide pairing) to intracellular trafficking from the ER to secretory granules and from exocytosis to target-cell signaling (33). A large literature pertains to sequence determinants of receptor binding and activation (67,68). Additional selective pressures presumably accompany distinctive properties of insulin not shared by IGFs (and vice versa): *e*.*g*., proteolytic processing of proinsulin to liberate the two-chain hormone and its self-assembly in the secretory granules of pancreatic β-cells (30) (and conversely IGF-specific binding to partner proteins; IGFBPs (69)). The invariant structural framework of insulin and IGFs provides a scaffold for specialization of cognate receptor-binding surfaces in the co-evolution of homologous receptor tyrosine kinases (65).

The mutant proinsulin syndrome has highlighted sequence determinants of nascent folding efficiency (52,70). Whereas classical comparison of species variants enabled study of divergent positions (71), advances in medical genetics has provided (and continues to provide) an ever-expanding database of *INS* mutations as unsuccessful “experiments of nature” (For recent review, see (27)). Such mutations generally affect conserved sites (47). Guided by clinical correlations, the present study thus focused on position B18, broadly conserved among vertebrate insulins as Val (but not invariant).^[4]^ Patients have been identified with mutations Ala^B18^ (DM onset in adolescence) (40) and Gly^B18^ (DM onset in infancy) (39). Because of this salient difference in ages of onset, we sought to investigate the relative impact of these substitutions on folding, structure, stability and function. Our approach combined chemical protein synthesis with cell-based interrogation of ER stress. Foreshortened single-chain insulin precursor polypeptides (“mini-proinsulins”) were prepared by SPPS using the “DesDi” framework established by DiMarchi and colleagues; in this 49-residue precursor Pro^B28^ is substituted by Lys and directly followed by Gly^A1^ (residues B29 and B30 are absent) (53). Folding of DesDi is more robust than longer precursors (including proinsulin) to unfavorable substitutions, presumably because the B28-A1 peptide bond facilitates formation of nascent structure that guides on-pathway disulfide pairing. In this framework Ala^B18^ and Gly^B18^ perturbed folding yields (relative to Val^B18^ DesDi) by ∼twofold and ∼8-fold, respectively. A similar pattern of relative perturbations to the thermodynamic stabilities of the respective single-chain analogs was also observed in CD-detected chemical denaturation experiments (Table 1). Although native or near-native α-helix contacts were preserved in the single-chain analogs, the guanidine-induced unfolding transition of the Gly^B18^ analog exhibited marginal cooperativity and a marked reduction in *m*-value (column 3 in table 1). Because the latter probes extent of exposure of nonpolar surfaces on denaturation, the low *m*-value of Gly^B18^ DesDi indicates aberrant pre-exposure of such surfaces in the folded state. These *in vitro* studies were extended through measurements of ER stress induced on transient transfection of WT or variant proinsulins in a human cell line: in these assays Gly^B18^ led to more robust induction of ER-stress signaling than Ala (Table S3). Ages of DM onset therefore correlate with the relative folding yields, native-state stabilities and degree of cellular ER stress.

To gain additional insight into structure and function, the single-chain synthetic precursors were cleaved by endo-Lys C to yield two-chain insulin analogs. The parent analog (*des*[B29, B30]-Lys^B28^-insulin) exhibits nativelike structure and function. Unlike CD spectra of the single-chain precursors, spectra of the two-chain analogs exhibited significant differences. Although Ala^B18^ was associated with a slight accentuation of α-helical features, this two-chain analog nonetheless exhibited impaired stability (ΔΔG_u_ 1.2(±0.2) kcal/mole). Strikingly, Gly^B18^ induced a twofold attenuation of α-helix content with complete loss of cooperativity; the partial fold is thus a molten globule (72). Whereas ^1^H- and ^13^C-NMR studies of the Ala^B18^ two-chain analog corroborated preservation of nativelike chemical shifts and inter-residue NOEs, spectra of the Gly^B18^ analog exhibited limited chemical-shift dispersion and attenuated inter-residue NOEs. A subset of long-range NOEs was nonetheless retained (as inferred based on presumptive assignments), suggesting that the Gly^B18^ molten globule is not a random coil but instead maintains on average some native-like features (73). We speculate that maintenance of the three canonical disulfide bridges enforces a subset of nativelike tertiary contacts, at least on average in a flexible ensemble of structures. The low but not negligible folding yield of the Gly^B18^ single-chain precursor is likely enabled by the B28-A1 peptide bond, which is in effect a fourth bridge that enforces native-like structural relationships (74). Cleavage of the B28-A1 peptide bond further relaxes this constraint leading to a molten partial fold. Because the 35-residue connecting domain of proinsulin (together with residues B29-B30) provides only a long flexible tether between the main-chain atoms of residues B28 and A1, we anticipate that (a) Gly^B18^-proinsulin would likewise form a molten globule and (b) the present DesDi-based synthetic studies of a Gly^B18^ single-chain precursor (however markedly perturbed) over-estimate the degree to which this mutation would block the oxidative folding of full-length proinsulin (75,76). It would be of future interest to evaluate the folding yields of Gly^B18^ single-chain precursor polypeptides (relative to corresponding Val^B18^ and Ala^B18^ analogs) as a function of connecting-domain length in the range 0-35 residues.

The mechanistic origins of the perturbations observed in the Ala^B18^ and Gly^B18^ analogs (relative to WT Val) may be considered in relation to general biophysical principles. Three properties may be pertinent: (a) relative hydrophobic transfer free energies; (77-79) (b) relative helical propensities (57,58,80) and (c) creation of a destabilizing cavity or crevice (81,82) created by the mutations as “large-to-small” substitutions (82). The first two properties reflect intrinsic features of these amino-acid types; the third depends also on structural details of the insulin molecule. This three-part analysis is given in Table 2. The mild perturbation introduced by Ala^B18^ is the net result of favorable and unfavorable terms. Whereas Ala exhibits a higher helical propensity than Val (as a β-branched side chain) (57,58,80), its hydrophobic transfer free energy is lower than that of Val (79). Because Val^B18^ is largely buried, net destabilization is predicted on account of Ala’s smaller volume, which leads to a cavity penalty (81,82). This cavity penalty is even larger for Gly^B18^ as is the hydrophobic transfer free-energy penalty; unlike in the Ala^B18^ analog, these unfavorable terms are not compensated or offset by a more favorable helix-propensity term. Our findings are thus in accordance with general models of protein stability (83). Global loss of helix content and organized structure in the Gly^B18^ two-chain molten globule is presumably driven by entropic terms, i.e., minimization of the entropic cost of folding to a native state whose putative stability (in the context of the WT structure) would suffer three independent perturbations (a-c).

### Concluding Remarks

The diverse mutations associated with the mutant insulin syndrome exhibit a broad range of ages of onset (84). The mildest, with onset in early adulthood, may even exhibit variable genetic penetrance (38,85); the most severe present in the neonatal period (24,25). The mutations typically exhibit genetic dominance such that expression of the variant proinsulin interferes with biosynthesis and secretion of WT proinsulin (33). Although such patients currently require insulin replacement therapy (86), it is possible that in the future such monogenic forms of DM may be treated through pharmacologic or gene-based therapies designed either to attenuate β-cell ER stress or to restore folding to the variant proinsulin (87).

We imagine that a panel of variant proinsulins (beyond Gly and Ala at position 18) might provide spectrum of relative foldabilities--in essence a calibrated “genetic rheostat” of ER stress levels--as a model for development of novel therapeutics: a case study in precision medicine (88). Given the general role of chronic ER stress in non-syndromic Type 2 DM (i.e., as induced by compensatory over-expression of WT proinsulin in the face of obesity-related insulin resistance), such δ-cell-directed therapies might address the growing pandemic of prediabetes leading to “diabesity” as a pending crisis in global health (89,90).

### Experimental procedures

Additional details pertaining to materials and methods are provided in the Supplement.

#### Chemical peptide synthesis

Peptides were synthesized on Tribute (Gyros Protein technologies) 2-Channel peptide synthesizer using 6-Cl-HOBt and DIC (1:1, equimolar with respect to Fmoc-protected amino acids) as the coupling agents and 20% piperidine in DMF as deprotection agent. After cleavage, peptide was precipitated in ether and dried.

#### Folding of DesDi mini-proinsulin precursor and its two-chain conversion

The crude 49-mer peptides, after the Fmoc-SPPS, was subjected to folding conditions (53): 0.1 mM peptide, 20 mM glycine, 2 mM cysteine at pH 10.5 for 16 hr.

Single-chain DesDi analogs were treated with Endo Lys-C enzyme (91) in 25 mM Tris base, 100 mM urea buffer (pH 8.5) at 12 °C water bath for 24 h. After HPLC indicated two chain conversion (typically 60-80%), the reaction mixture was acidified and purified on semi-preparative HPLC. Fractions containing clean protein were pooled and lyophilized.

### Chain Combination

Wild-type A- and B chains were obtained by sulfitolysis of commercially available human insulin. Variant B chains were made by solid-phase peptide synthesis and converted to their respective S-sulfonate derivatives. For chain-combination reactions, a stock solution of WT A-chain S-sulfonates was first prepared and split three ways before mixing with respective B-chain S-sulfonate analogs (WT, Ala^B18^ or Gly^B18^); the reaction proceeded in 0.1 M glycine buffer at pH 10.5 using 1 equiv. of dithiothreitol and stirred at 4 °C for 2 days; details are provided in the Supplement.

#### Circular Dichroism (CD)

Samples were quantitated using Genesis 150 UV-Vis spectrophotometer (Thermo Scientific) using one-cm cuvette by measuring peptide absorbances at 280 nm. The extinction coefficient used was: 6335 M^-1^ cm^-1^.

Far-ultraviolet (UV) spectra were obtained on a Jasco J-1500 spectropolarimeter equipped with an automated syringe-driven titration unit. Spectra were obtained from 195-255 nm as described (92). Insulin analogues were made 50 μM in a buffer containing 10 mM potassium phosphate (pH 7.4) and 50 mM KCl. Helix-sensitive wavelength 222 nm was used as a probe of denaturation of protein (at 5 μM) by guanidine hydrochloride. Thermodynamic parameters were obtained by application of a two-state model as described (93).

#### NMR spectroscopy

^1^H NMR spectra of DesDi two chain analogs were acquired at a proton frequency of 700 MHz at pD 7.4 (direct meter reading) at 32 °C. ^1^H-^13^C heteronuclear single-quantum coherence (HSQC) spectra were acquired at natural abundance as described. The spectra were obtained at ^13^C frequency of 176 MHz at constant temperature of 305 K using the “hsqcetgp” Bruker pulse sequence as described by the vendor. Acquisition with FID size 2048 × 128, 800 scans, 1.0 sec relaxation delay, sweep widths 11 ppm (^1^H) and 70 ppm (^13^C) with offset 4.7 and 40 ppm for the ^1^H and ^13^C dimension, respectively. Data were processed with Topspin 4.0.6 (Bruker Biospin) and analyzed with Sparky software (94) using a 90° shifted-sine window function to a total of 2048 × 1024 data points (F2 × F1), followed by automated baseline- and phase correction. All NMR data were acquired using a BRUKER 700 MHz spectrometer equipped with ^1^H, ^19^F, ^13^C, ^15^N quadruple resonance cryoprobe.

#### Cell culture

Human embryonic kidney 293T cell was cultured in Dulbecco’s Modified Eagle Medium (DMEM), supplemented with 10% fetal bovine serum (FBS), 1% penicillin/streptomycin as recommended by the American Type Culture Collection. Human hepatocellular carcinoma cell line HepG2 was also cultured in DMEM with 10% FBS and 1% penicillin/streptomycin. The protocol of signaling assays, which employed 24-h serum starvation, was adapted from previously published protocol (23). After starvation, cells were treated in parallel with a set of insulin analogs in serum-free medium.

#### Transient transfections and ER stress assays

Transfections were performed using Lipofectamine 3000 as described by the vendor (Invitrogen). Transfected HEK 293T cells were subjected to the Bio-Rad one-step real-time qPCR protocol. Readouts were provided by the up-regulation of ER stress markers *CHOP* and *BiP*. mRNA (messenger ribonucleic acid) abundances were measured in triplicate. In Western blot assay probing ER stress markers (60,95), after 24 hour post transient transfection, cells were lysed by RIPA buffer (Cell Signaling Technology; CST). Protein concentrations in lysates were measured by BCA assay (Thermo)- and subjected to 4-20% SDS-PAGE and WB using anti-pPERK. Anti-PERK, anti-BiP and anti-CHOP antibodies (CST) at a dilution ratio of 1:1000; GAPDH provided a loading control.

#### In-cell pIR immunoblotting

The insulin-dependent IR activation was probed via fluorescent readouts. HepG2 cells were seeded (∼8000 cells/well) into a 96-well black plate. After serum starvation, serial analog dilutions (100 μl) the following treatments were applied to each well as described (23). Fixed cells were then exposed to the primary antibody (10 μL anti-pTyr 4G10) and the secondary antibody (anti-mouse-IgG-800-CW antibody (Sigma) in 25 ml Blocking Buffer) was added to enable measurement of the fluorescence signals (details in supplemental information and ref (23)).

#### Cell-based insulin signaling assays

HepG2 cell line was subjected to 24-h serum-free starving, followed by addition of medium containing testing analogs (50 nM for ordinary treatment, additional 3000 nM for Ala^B18^ and Gly^B18^ DesDi two chain analogs). After lyzed by RIPA buffer, total protein concentrations of lysate were determined by BCA assay (Thermo). Blotting protocols were modified from previous publication (23) (for detail, see the supplemental information).

#### Rat Experiments

Animals were maintained in accredited facility of Case Western Reserve University School of Medicine. All procedures were approved by the Institutional Animal Care and Use Committee (IACUC) office of the University. Animal care and use was monitored by the University’s Veterinary Services.

#### Measurement of the Glucose-Lowering Effect of Insulins in Diabetic Rats

Male Lewis rats (average body mass of ∼300 g) were rendered diabetic by streptozotocin (STZ), as previously described (96). DesDi two chain insulins were dissolved in Lilly® Diluent buffer with the specified dose and injected in 100 μL/300 g rat. Control rats received the appropriate volume of the Lilly buffer. For these subcutaneous (SQ) experiments, rats were injected under the skin into the soft tissue in the posterior aspect of the neck. Following injection, blood glucose was measured using a small drop of blood obtained from the clipped tip of the rat’s tail using a clinical glucometer (EasyMax® V Glucose Meter, Oak Tree Health, Las Vegas, NV). Blood-glucose concentrations were measured at t=0, and every 10 min for the first hour (h), every 20 min for the second h, every 30 min for the third h, and then each h for the rest of the experiment.

### Statistical Analysis

Data analysis was performed using MATHLAB Online (MathWorks). Areas under the curve (AUC) of glucose values in the diluent- and insulin-injected rats were calculated (Fig. S20). Statistical significance was determined using ANOVA; a *p* value of <0.05 was considered significant whereas *p* values just above this threshold may suggest a trend whose validation would require larger studies.

## Acknowledgments

This work was supported in part by the National Institutes of Health Grant (R01 DK040949 to M.A.W.). Early stages of this work were also supported by a grant from the NIH to M.A.W. and P.A. (R01 DK069764) and to P.A. (R01 DK48280). The content is solely the responsibility of the authors and does not necessarily represent the official views of the National Institutes of Health. Y.-S.C. was supported in part by a foundation grant from the Diabetes Research Connection. N.F.B.P. was supported in part by the American Diabetes Association Grants 7-13-IN-31 and 1-08-RA-149. We M.C. Lawrence and N. Rege for helpful discussion; and G.I. Bell and L.H. Philipson (University of Chicago) for summary of clinical mutations in the human insulin gene. M.A.W. thanks M. Karplus (Harvard), T. Sosnick (University of Chicago) and the late P.G. Katsoyannis (Mt. Sinai Medical Center, NY, NY) for helpful discussion in the early stages of this work. We dedicate this article to the memory of the late Prof. G.G. Dodson (University of York, York, UK).

## Author Contributions

B.D. and M.J. undertook chemical peptide syntheses and folding assays; B.D. performed CD studies; Y.S.C. performed cell-based assays; Y.S.C., and M.A.W. designed and interpreted functional assays; Y.Y. and M.A.W. designed, performed and interpreted NMR studies; B.D., D.C. and M.A.W. interpreted cryo-EM structures of insulin-receptor model complexes; R.G., N.F.B.P. and F.I.-B performed rat studies; B.D., Y.S.C., Y.Y., D.C., and M.A.W. prepared figures; B.D. and M.A.W. conceptualized the program of research. All authors contributed commented on the manuscript.

## FOOTNOTES

## ^1^Abbreviations

AKT: protein kinase B
AUC: areas under the curve in insulin tolerance tests
BiP: binding immunoglobulin protein
CHOP: C/EBP homologous protein
CD: circular dichroism
CFTR: cystic fibrosis transmembrane conductance regulator
ER: endoplasmic reticulum
DM: diabetes mellitus
GRP: glucose-regulated proteins
HEK: human embryonic kidney cell line
HepG2: human liver cancer cell line
HPLC: high-performance liquid chromatography
HSQC: heteronuclear single quantum coherence
IGF: insulin-like growth factor
IGFBP: insulin-like growth factor binding protein
SPPS: solid-phase peptide synthesis
ISR: integrated stress response
IR: insulin receptor
NMR: nuclear magnetic resonance
NOE: nuclear Overhauser effect
NOESY: NOE spectroscopy
PDB: Protein Data Bank
PERK: protein kinase R-like ER kinase
qPCR: quantitative polymerase chain reaction
STZ: streptozotocin
SQ: subcutaneous
TK: tyrosine kinase
TOCSY: total correlation spectroscopy
WT: wild-type. WT represents human insulin whereas the “ parent” DesDi precursor is Lys^B28^-*des*-dipeptide[B29,B30]-mini-proinsulin. Amino acids are designated by standard one- or three-letter codes.

## Footnotes

2 The mutant proinsulin syndrome can thus present as permanent neonatal DM or as a subtype of the clinical syndrome *m*aturity-*o*nset *d*iabetes of the *y*oung (MODY) (97); this spectrum of monogenic DM is collectively designated “<u>m</u>utant *ins* gene-induced <u>d</u>iabetes of <u>y</u>outh” (MIDY) (26,51).

3 Lys^B28^ is a substitution in insulin *lispro*, the active component of Humalog^®^, a prandial analog formulation (Eli Lilly and Co.) (98). In broad clinical use, this analog contains paired substitutions in the B chain (Pro^B28^→Lys and Lys^B29^→Pro) that destabilize the zinc insulin hexamer (99) leading to accelerated SQ absorption (100).

4 Rare species variants include related branched-chain residues Ile (in some snakes) and Thr in the amyloidogenic insulin of an hystricomorph rodent (the degu rat; Supplemental Fig. S1); Ile is also observed at position B18 of relaxin-related members of the vertebrate insulin superfamily (Supplemental Fig. S2).

