## Supplemental Information for "Age of diabetes onset in the mutant proinsulin syndrome correlates with mutational impairment of protein foldability and stability"

### Purpose of Supplement

Fourteen Supplemental Figures and three Supplemental Tables are provided. These give additional data and details regarding methods, protein sequences, structural models, chemical polypeptide syntheses, cell-based ER stress assays and STZ diabetic rat studies.

Figure S16. Far-UV CD temperature/wavelength spectra……………………….…..………...15
Figure S17. Nature abundance $^{1}$H-$^{13}$C HSQC NMR spectra ……….…………………..……....16
Figure S18. Temperature-dependent $^{1}$H-NMR studies …….……………………..………....17
Figure S19. Concentration- and temperature-dependent NMR spectra of DesDi-Ala$^{B18}$……......18
Figure S20. Areas under the curve (AUC) analysis of rat studies ………….........................19
Table S1. Isolated yields from folding of DesDi single-chain analogs …...................................20
Table S2. CD-derived helical parameters ……………..............................................................20
Table S3. Responses of ER stress markers to Ala$^{B18}$ and Gly$^{B18}$ in human proinsulin …............21
Table S4. Side-chain distances (Å) between residue B18 and neighboring residues ..................21
Supplemental References ………………..................................................................22

**Experimental Procedures**: **Additional Details**

*Materials*—Fmoc amino acids were purchased from Gyros Protein Technologies. Protecting groups were: Arg(Pbf), Asn(Trt), Cys(Trt), Gln(Trt), Glu(OtBu), His(Trt), Lys(Boc), Ser(tBu), Thr(tBu), Tyr(tBu). H-Asn(Trt)-HMPB-CM resin (0.44 mmol/g loading), 6-chloro-1-hydroxybenzotriazole (6-Cl-HOBt) and *N,N′*-diisopropylcarbodiimide (DIC) were obtained from Gyros Protein Technologies. Piperidine and trifluoroacetic acid were purchased from Chem-Impex International. Diethyl ether, dichloromethane (DCM), *N,N*-dimethylformamide (DMF), acetonitrile (HPLC-grade), and guanidine hydrochloride (GuHCl) were purchased from Fisher. All other reagents were purchased from Sigma-Aldrich and were of the purest grade available.

*Analytical HPLC conditions*—Analytical reverse-phase high-performance liquid chromatography (HPLC) was performed using a Waters 1525 Binary HPLC system. All chromatographic separations were performed on a C8 Proto (4.6 mm x 250 mm) 300-Å, 5-µm, Higgins Analytical Inc. column, using 25-50% Solvent B in Solvent A over 35 minutes (min). (solvent A = 0.1% trifluoroacetic acid (TFA) in HPLC-grade de-ionized, distilled water; solvent B = 0.1% TFA in HPLC-grade acetonitrile) at a flow rate of 1.0 mL/min with detection by UV absorption at 215 nm.

*Preparative HPLC conditions*—A Waters 2545 Quaternary pumping system (equipped with FlexInject) was used for preparative HPLC purifications. Chromatographic separations were performed on a C4 Proto (20 mm x 250 mm) 300-Å, 10-µm Higgins Analytical Inc. column, using 25-50% Solvent B in Solvent A (ass above) over 35 min at a flow rate of 20 mL/min with detection by UV absorption at 215 nm. Fractions containing the desired products were identified by analytical LC and mass spectrometry, then combined and lyophilized.

*LC-MS*—Liquid chromatography-mass spectrometry (LC-MS) analysis was performed on LCQ Advantage Ion Trap Mass Spectrometer System coupled to an Agilent 1100 Series HPLC system. Masses were obtained by online electrospray mass spectrometry. MS data shown were collected across the entire principal UV-absorbing peak in each chromatogram.

*Peptide synthesis*—DesDi 49-mer peptides were synthesized on Tribute (Gyros Protein technologies) 2-Channel peptide synthesizer. All the amino acids were 10-times excess relative to the resin. 6-Cl-HOBt and DIC (1:1, equimolar with respect to Fmoc-protected amino acids) were used as the coupling agents and 20% piperidine in DMF used for deprotection. At the end of the synthesis, peptides were cleaved with Trifluoroacetic acid (TFA) cocktail containing 2.5% *v/v* of 2,2′-(ethylenedioxy)diethanethiol, triisopropylsilane, anisole and water. Cleavage mixture was precipitated with ether (5-10 fold with respect to TFA) and solid was isolated by centrifugation. Precipitate was further washed with twice with ether and dried in vacuo.

*Folding of DesDi-precursor and two chain conversion*—The crude 49-mer peptides, after the Fmoc-SPPS, was subjected to folding conditions following published protocol (1): 0.1 mM peptide, 20 mM glycine, 2 mM cysteine at pH 10.5 for 16 hr. After HPLC indicated the sharp early eluting peak, the reaction mixture acidified to pH 3.0, filtered (0.22 µ) and purified on a reverse phase (RP) C4-preparative column.

Single chain DesDi analogs at 4mg/mL concentration were treated with Endo Lys-C enzyme (2) in 25 mM Tris base, 100 mM urea buffer (pH 8.5). The reaction mixture was kept at 12 °C water bath for 24 h. After HPLC indicated two chain conversion (typically 60-80%), the reaction mixture was acidified to pH 2-3 and purified on semi-preparative HPLC using 25-50% Solvent B in Solvent A over 35 minutes. Fractions containing clean protein were pooled and lyophilized.

*Chain Combination reaction*—To get unmodified A- and B- chains for the chain combination, wild type insulin (75 mg) was subjected to sulfitolysis reaction in a mixture containing $Na_2S_2O_4$ (65 mg) and $Na_2SO_3$ (130 mg) dissolved in 4 mL buffer (5 M GuHCl, 50 mM phosphate, pH 7.0). Reaction stirred for 3 hours at room temperature and directly purified on C18 Triart column using $NH_4HCO_3$ (25 mM, pH 8.0)/ acetonitrile gradient 25-50%. This reaction provided 35 mg of A-chain S-Sulfonate and 42 mg of B-chain S-sulfonate. Purity of the materials was confirmed by HPLC and mass was confirmed by LCMS.

$Ala^{B18}$ and $Gly^{B18}$ modified B-chains were first synthesized by solid phase peptide synthesis. Purified peptides were subjected to sulfitolysis reaction conditions as described above. After stirring for 3 hours at room temperature, the reaction was purified to get the respective B-chain S-Sulfonates. The purity of S-sulfonates was confirmed by rp-HPLC and mass was confirmed by LCMS (Fig. S12)

Stock solutions of 2.4 mM insulin A-chain S-sulfonate (5 mg/0.8 mL), 0.8 mM WT B-chain S-sulfonate (0.85 mg/0.25 mL), 0.8 mM $Ala^{B18}$ B-chain S-sulfonate (0.85 mg/0.25 mL) and 0.8 mM $Gly^{B18}$ B-chain S-sulfonate (0.85 mg/0.25 mL) were prepared in 0.1 M glycine buffer pH 10.). Three reactions were set up by mixing WT A-chain S-sulfonate stock (0.25 mL) with each of the B-chain S-sulfonate and adding a 1.0 equiv. of DTT. The reactions were monitored over 2 days using LCMS (Fig. S13-15).

*Transient transfections and ER stress assays*—Transfections were performed using Lipofectamine 3000 as described by the vendor (Invitrogen). After 8 hours (h) in Opti-MEM medium, cells were recovered using fresh culture medium with fetal bovine serum (FBS). Following transient transfection, HEK 293T cells were subjected to the Bio-Rad one-step real-time qPCR protocol. Readouts were provided by the up-regulation of ER stress markers *CHOP* and *BiP*. mRNA (messenger ribonucleic acid) abundances were measured in triplicate by quantitative polymerase chain reaction. Samples were prepared as described by the vendor (One-Step rt-PCR reagent kits; Bio-rad). The primers were used essentially as described (3,4). In ER-stress-relataed Western blot (WB) assays, after 24 h post-transient transfection, cells were lysed by RIPA buffer (Cell Signaling Technology; CST). Protein concentrations were measured by BCA assay (Thermo); cell lysates were subjected to 4-20% SDS-PAGE and WB using anti-pPERK, anti-PERK (5), anti-BiP and anti-CHOP antibodies (CST) at a dilution ratio of 1:1000; GAPDH provided a loading control. Western blot used 4-20% SDS-PAGE and revealed by HRP-conjugated secondary antibody (CST).

*In-cell pIR immunoblotting*—The insulin-dependent IR activation was probed via fluorescent readouts. HepG2 cells were seeded (~8000 cells/well) into a 96-well black plate. After serum starvation in 100-µl plain Hanks' Balanced Salt Solution (HBSS) for 2 h at 37 °C, serial analog dilutions (100 µl) were applied to each well; cells were then incubated for 20 min at 37 °C, followed by the 3.7% formaldehyde fixation. 200 µl of 0.1% Triton-X-100 (Sigma) was then added to permeabilize the cells, followed by their fixation by 100-µl Odyssey Blocking Buffer (LI-COR). A blocking procedure was applied for 1 hr at room temperature on an orbital shaker. Fixed cells were then exposed to the primary antibody (10 µL anti-pTyr 4G10 into 20 ml Blocking Buffer) overnight at 4 °C. The secondary antibody (anti-mouse-IgG-800-CW antibody (Sigma) in 25 ml Blocking Buffer) was added after a wash. pTyr was detected via 800 nm emission. DRAQ5 (Fisher) was also applied to enable measurement of cell number via 700 nm emission. The fluorescence signals were detected on a LI-COR Infrared Imaging system (Odyssey) under settings as follows: offset 4 mm with setting "Intensity-Auto" (6).

*Signaling assays in a mammalian cell line*—HepG2 cell line was subjected to 24-h serum-free starving, followed by addition of medium containing testing analogs (50 nM for ordinary treatment, additional 3000 nM for $Ala^{B18}$ and $Gly^{B18}$ DesDi two chain analogs). After lyzed by RIPA buffer containing protease and phosphatase inhibitor cocktails (Roche), total protein concentrations of lysate in RIPA buffer were determined by BCA assay (Thermo). Blotting protocols were modified from previous publication. Briefly, for p-IR/IR blotting, samples were probed by insulin receptor β (4B8) antibody (CST antibodies unless otherwise stated) or an equal mixture of anti-phospho-insulin receptor β (Tyr1150/1151); phospho-insulin receptor (Tyr1158) antibody (Thermo); phospho-insulin receptor (Tyr1334) antibody (Thermo); phospho-insulin receptor β (Tyr1345) mAb; and anti-phosphor-insulin receptor (phospho-Tyr972) antibody (Abcam). Dilutions for these antibodies were 1:5000 in 5% bovine serum albumin. Antibodies for Akt blotting were p-Akt antibody (Ser473) (1:1000) and Akt1/2/3 antibody (H-136) (1:1000) (6).

| | B chain (1 … 15, 18 … 30) | A chain (1 … 13, 16 … 21) |
|---|---|---|
| **mammals** | | |
| *human* | FVNQHLCGSHLVEALYLVCGERGFFYTPKT | GIVEQCCTSICSLYQLENYCN |
| *dog* | FVNQHLCGSHLVEALYLVCGERGFFYTPKA | GIVEQCCTSICSLYQLENYCN |
| *cat* | FVNQHLCGSHLVEALYLVCGERGFFYTPKA | GIVEQCCASVCSLYQLEHYCN |
| *pig* | FVNQHLCGSHLVEALYLVCGERGFFYTPKA | GIVEQCCTSICSLYQLENYCN |
| *cattle* | FVNQHLCGSHLVEALYLVCGERGFFYTPKA | GIVEQCCASVCSLYQLENYCN |
| *sheep* | FVNQHLCGSHLVEALYLVCGERGFFYTPKA | GIVEQCCAGVCSLYQLENYCN |
| *beluga Whale* | FVNQHLCGSHLVEALYLVCGERGFFYTPKA | GIVEQCCTSICSLYQLENYCN |
| *rabbit* | FVNQHLCGSHLVEALYLVCGERGFFYTPKS | GIVEQCCTSICSLYQLENYCN |
| *gorilla* | FVNQHLCGSHLVEALYLVCGERGFFYTPKT | GIVEQCCTSICSLYQLENYCN |
| *chimpanzee* | FVNQHLCGSHLVEALYLVCGERGFFYTPKT | GIVEQCCTSICSLYQLENYCN |
| *bornean orangutan* | FVNQHLCGSHLVEALYLVCGERGFFYTPKT | GIVEQCCTSICSLYQLENYCN |
| *green monkey* | FVNQHLCGSHLVEALYLVCGERGFFYTPKT | GIVEQCCTSICSLYQLENYCN |
| **rodents** | | |
| *rat* | FVKQHLCGPHLVEALYLVCGERGFFYTPKS | GIVDQCCTSICSLYQLENYCN |
| *octodon Degu* | YSSQHLCGSNLVEALYMTCGRSGFYRPHD | GIVDQCCNNICTFNQLQNYCNVP |
| *guinea pig* | FVSRHLCGSNLVETLYSVCQDDGFFYIPKD | GIVDQCCTGTCTRHQLQSYCN |
| *porcupine* | FVNQHLCGSHLVEALYLVCGNDGFFYRPKA | GIVDQCCTGVCSLYQLQNYCN |
| *chinese hamster* | FVNQHLCGSHLVEALYLVCGERGFFYTPKS | GIVDQCCTSICSLYQLENYCN |
| **reptiles and amphibians** | | |
| *chameleon* | LPNQHLCGSHLVEALYLVCGDRGFYYSPKT | GIVQQCCENTCSLYELENYCN |
| *african clawed frog* | LVNQHLCGSHLVEALYLVCGDRGFFYYPKV | GIVEQCCHSTCSLFQLESYCN |
| *mainland tiger snake* | APNQRLCGSHLVEALFLICGERGFYYSPRE | GIVEQCCENTCSLYELENYCN |
| *central bearded dragon* | IPNQHLCGSHLVEALYLVCGERGFYYSPKT | GIVQQCCENTCSLYELENYCN |
| *alligator* | AANQRLCGSHLVDALYLVCGERGFFYSPKG | GIVEQCCHNTCSLYQLENYCN |
| **birds** | | |
| *chicken* | AANQHLCGSHLVEALYLVCGERGFFYSPKA | GIVEQCCHNTCSLYQLENYCN |
| *turkey vulture* | VANQHLCGSHLVEALYLVCGERGFFYSPKA | GIVEQCCHNTCSLYQLENYCN |
| *common ostrich* | AANQHLCGSHLVEALYLVCGERGFFYSPKA | GIVEQCCHNTCSLYQLENYCN |
| *Rufous hummingbird* | AVNQHLCGSHLVEALYLVCGERGFFYSPKA | GIVEQCCHNTCSLYQLENYCN |
| *Wild turkey* | AANQHLCGSHLVEALYLVCGERGFFYSPKA | GIVEQCCHNTCSLYQLENYCN |
| *brown roatelo* | AANQHLCGSHLVEALYLVCGERGFFYSPKA | GIVEQCCHNTCSLYQLENYCN |
| **fish** | | |
| *zebrafish* | GTPQHLCGSHLVDALYLVCGPTGFFYNPKR | GIVEQCCHKPCSIFELQNYCN |
| *common carp* | GAPQHLCGSHLVDALYLVCGPTGFFYNP | GIVEQCCHKPCSIFELQNYCN |
| *anglerfish* | APAQHLCGSHLVDALYLVCGDRGFFYNP | GIVEQCCHRPCNIFDLQNYCN |
| *barfin flounder* | LPPQHLCGAHLVDALYLVCGERGFFYTP | GIVEQCCHKPCNIFDLQNYCN |
| *Nile tilapia* | GGPQHLCGSHLVDALYLVCGDRGFFYNPR | GIVEECCHKPCTIFDLQNYCN |
| *butterflyfish* | ASSQHLCGSHLVDALYMVCGEKGFFYQPKT | GIVEQCCHHPCNIFDLQNYCN |
| *Chum salmon* | AAAQHLCGSHLVDALYLVCGEKGFFYTP | GIVEQCCHKPCNIFDLQNYCN |
| *ghost shark* | VPTQRLCGSHLVDALYFVCGERGFFYSPKQ | GIVEQCCHNTCSLVNLEGYCN |
| *spotted ratfish* | VPTQRLCGSHLVDALYFVCGERGFFYSPKPI | GIVEQCCHNTCSLANLEGYCN |
| *lamprey* | AGGTHLCGSHLVEALYVVCGDRGFFYTPSK | GIVEQCCHRKCSIYDMENYCN |
| *hagfish* | RTTGHLCGKDLVNALYIACGVRGFFYDPTK | GIVEQCCHKRCSIYDLENYCN |

**Figure S1**. Alignment of vertebrate insulin sequences (7). Motif-specific cysteines are shown in bold, and residue B18 in red within a red box. $Leu^{B15}$, $Leu^{A13}$ and $Leu^{A16}$ are highlighted in blue within blue boxes. $Cys^{B19}$ and $Cys^{A20}$ are shown within black box with yellow fill. Chain identifier (B or A) and residue numbers are indicated at top.

| | B chain (1 … 15 … 18 … 30) | A chain (1 … 13 … 16 … 21) |
|---|---|---|
| human ins | FVNQHLCGSHLVEALYLVCGERGFFYTPKT | GIVEQCCTSICSLYQLENYCN |
| IGF-1 | GPETLCGAELVDALQFVCGDRGFYFNKPT | GIVDECCFRSCDLRRLEMYCA |
| IGF2 | PSETLCGGELVDTLQFVCGDRGFYFSRPA | GIVEECCFRSCDLALLETYCA |
| REL1 | DDVIKLCGRELVRAQIAICGMSTWS | ALFEKCCLIGCTKRSLAKYC |
| REL2 | EEVIKLCGRELVRAQIAICGMSTWS | ALANKCCHVGCTKRSLARFC |
| INSL3 | EMREKLCGHHFVRALVRVCGGPRWSTEA | NPARYCCLSGCTQQDLLTLCPY |
| INSL4 | AAELRGCGPRFGKHLLSYCPMPEKTFTTTPGGWL | RFDPFCCEVICDDGTSVKLCT |
| INSL5 | KESVRLCGLEYIRTVIYICASSRW | DLQTLCCTDGCSMTDLSALC |
| INSL6 | SSARKLCGRYLVKEIEKLCGHANWSQF | GYSEKCCLTGCTKEELSIACLPY |

**Figure S2**. Sequence alignment of vertebrate insulin superfamily (8). The color code is the same as in Supplemental Figure S1.

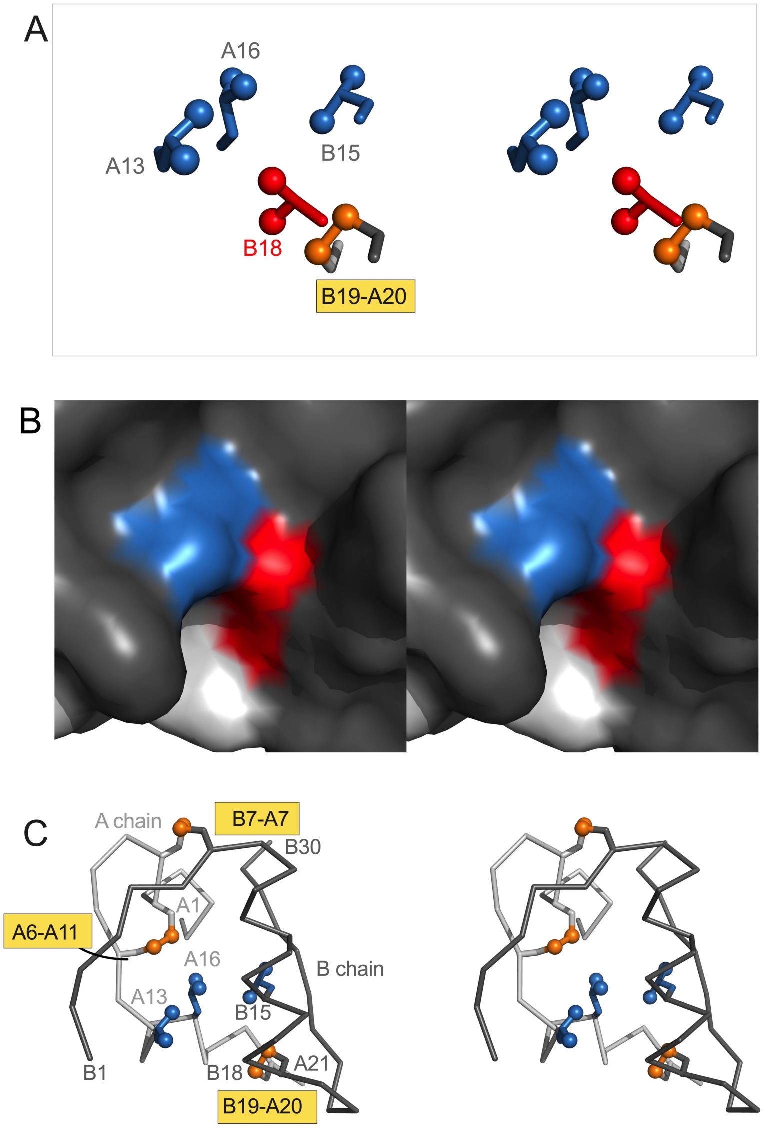

**Figure S3**. Spatial environment of $Val^{B18}$ in the insulin monomer (9). (A) Stereo view of $Val^{B18}$ and other neighboring side chains (as extracted from 2-Zn insulin hexamer; PDB entry 4INS (10)). Side chains of $Leu^{B15}$ (blue), $Val^{B18}$ (red), $Leu^{A13}$ (blue), $Leu^{A16}$ (blue) and cystines (golden) are shown as sticks. In Table S4 are given distances between side chains of $Val^{B18}$ (cutoff <6Å) and predicted distances to $Ala^{B18}$ (distances up to 8 Å) to their respective neighboring residues. (B) Protein surface model of the canonical insulin monomer in same orientation; the color code is as in panel A (A- and B chain surfaces are shown in light and dark gray, respectively). (C) Stereo view of $Gly^{B18}$ model with $Leu^{B15}$, $Leu^{A13}$ and $Leu^{A16}$ side chains in blue and disulfides in gold. Terminal methyl groups of these side chains and sulfur atoms in panel A and C are shown as spheres (one-third Van der Waals radii).

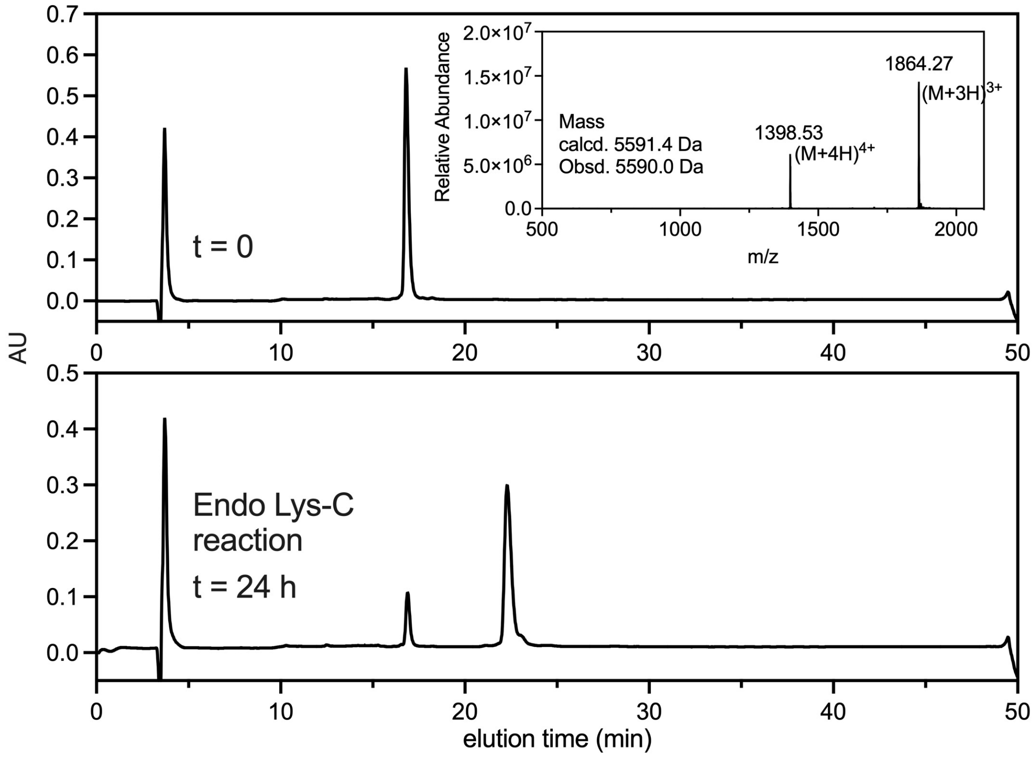

**Figure S4**. Analytical HPLC profiles for Endo-Lys-C digestion of Val$^{B18}$ (WT)-DesDi single-chain insulin (49 residues). Mass spectrum of single-chain analog is shown in inset.

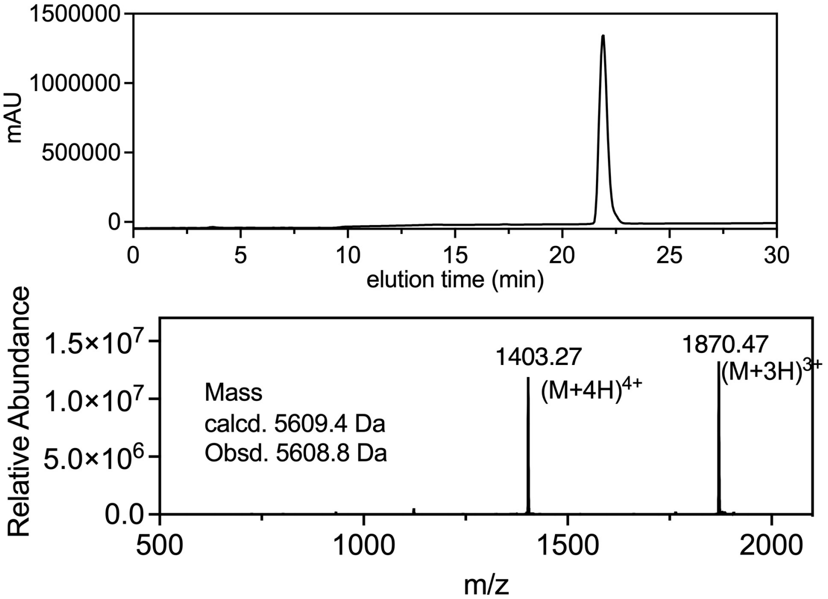

**Figure S5**. LC-MS data for the Val$^{B18}$ (WT)-DesDi two-chain insulin analog.

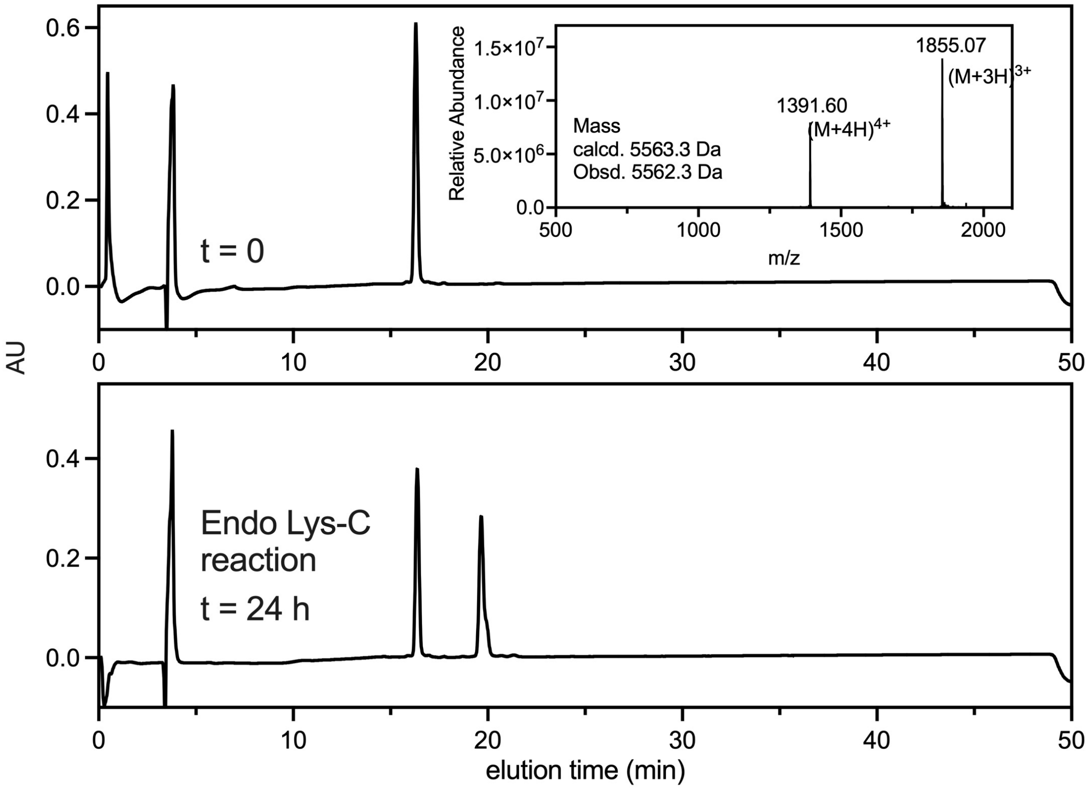

**Figure S6**. Analytical HPLC profiles for Endo-Lys-C digestion of Ala$^{B18}$-DesDi single-chain insulin. Mass spectrum is shown in inset.

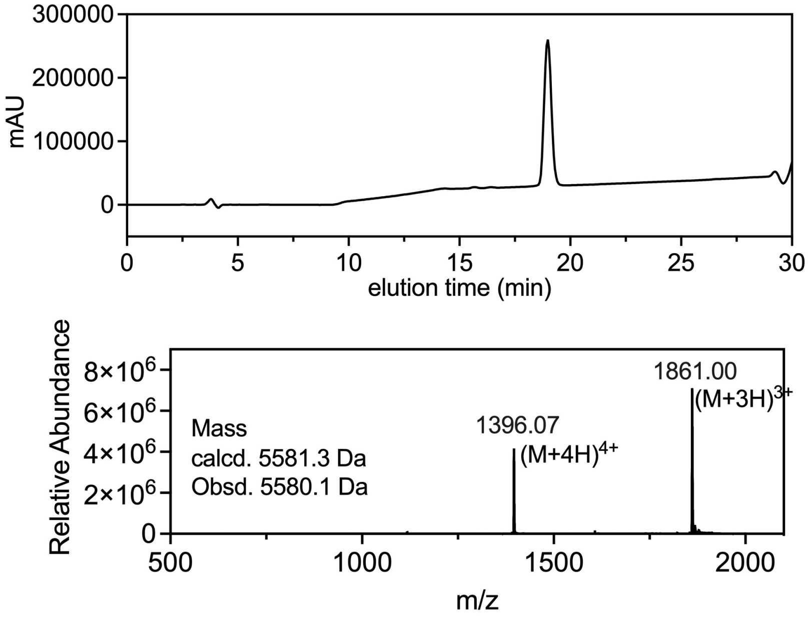

**Figure S7**. LC-MS data for the Ala$^{B18}$-DesDi two-chain insulin analog.

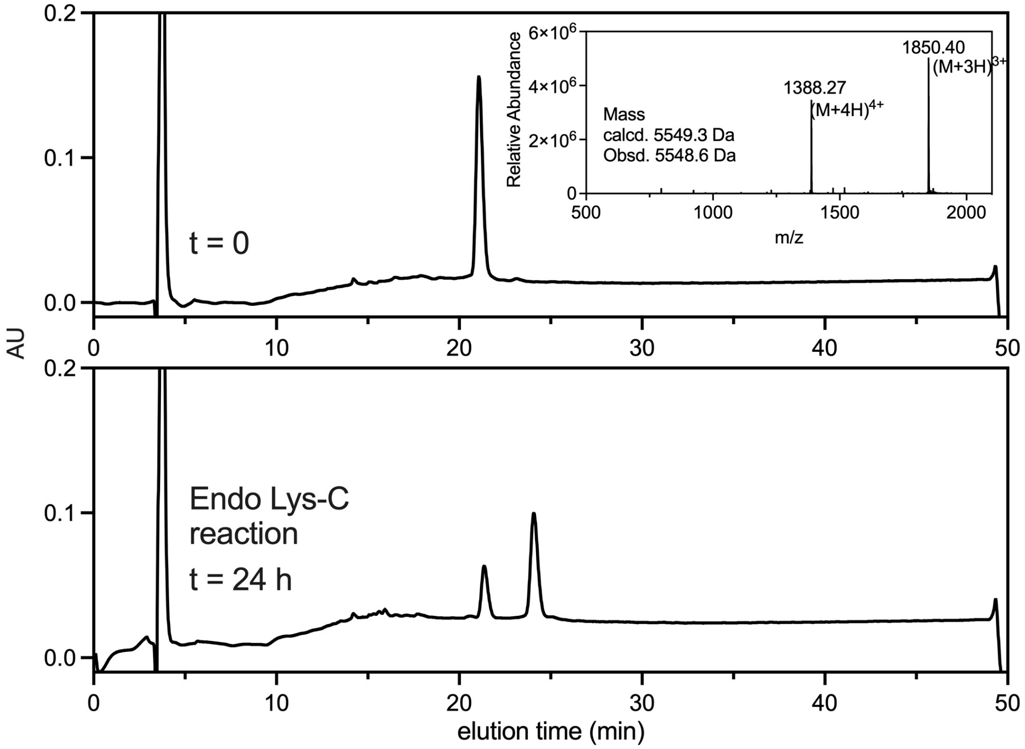

**Figure S8**. Analytical HPLC profiles for Endo-Lys-C digestion of Gly$^{B18}$-DesDi single-chain insulin analog. Mass spectrum is shown in inset.

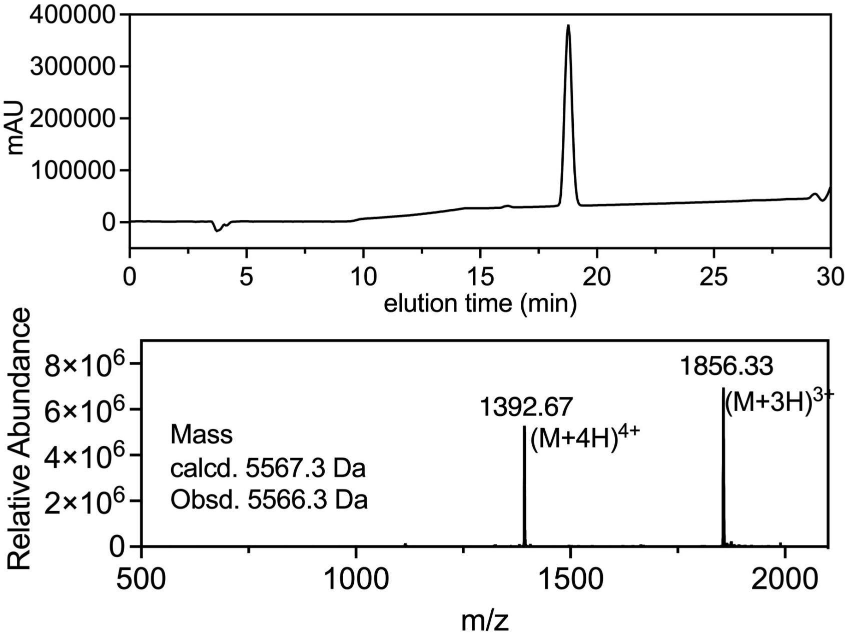

**Figure S9**. LC-MS data for the Gly$^{B18}$-DesDi two-chain insulin.

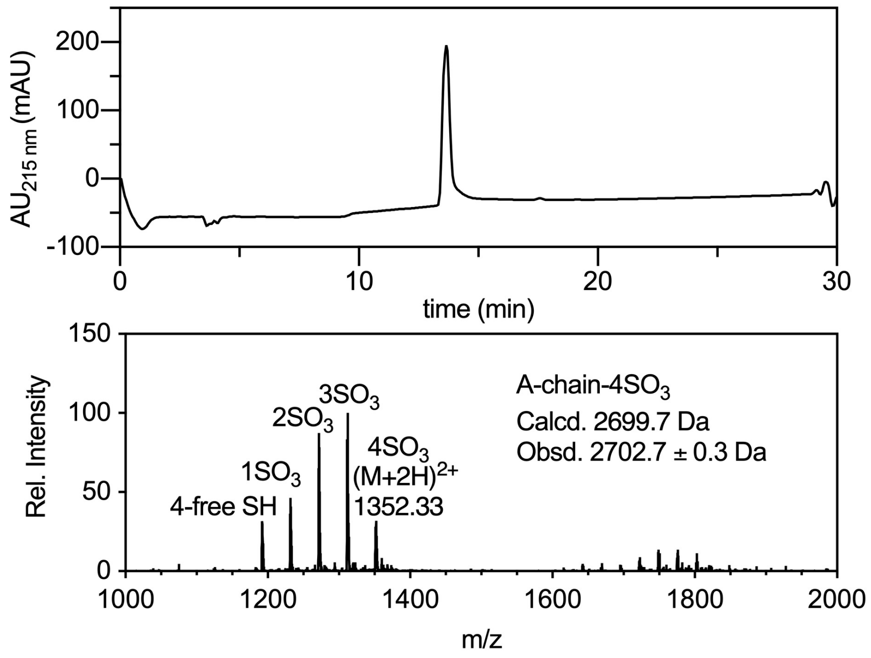

**Figure S10.** LCMS analysis of isolated insulin A-chain (S-$SO_3$) peptide.

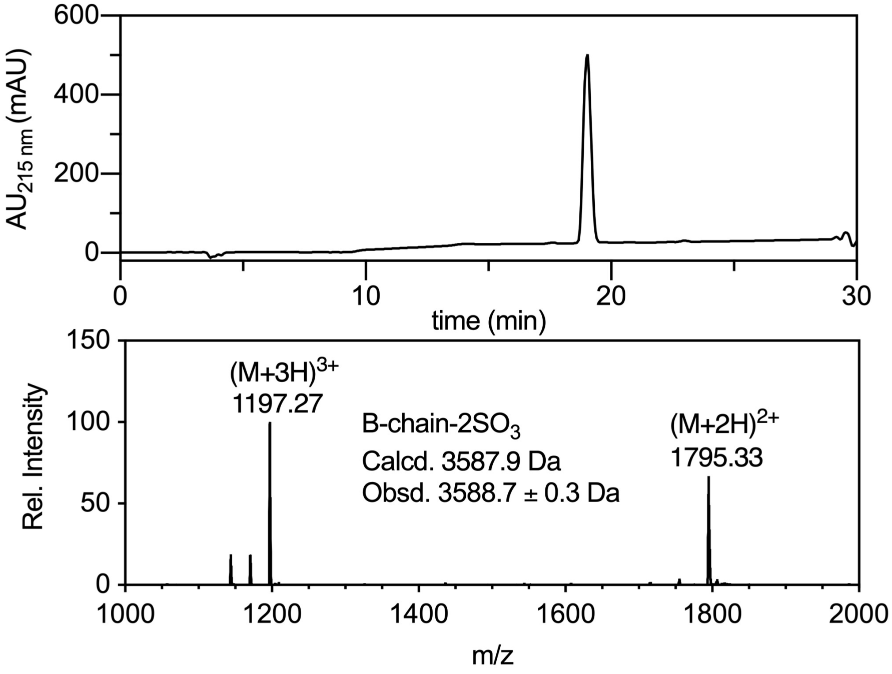

**Figure S11.** LCMS analysis of isolated insulin B-chain (S-$SO_3$) peptide.

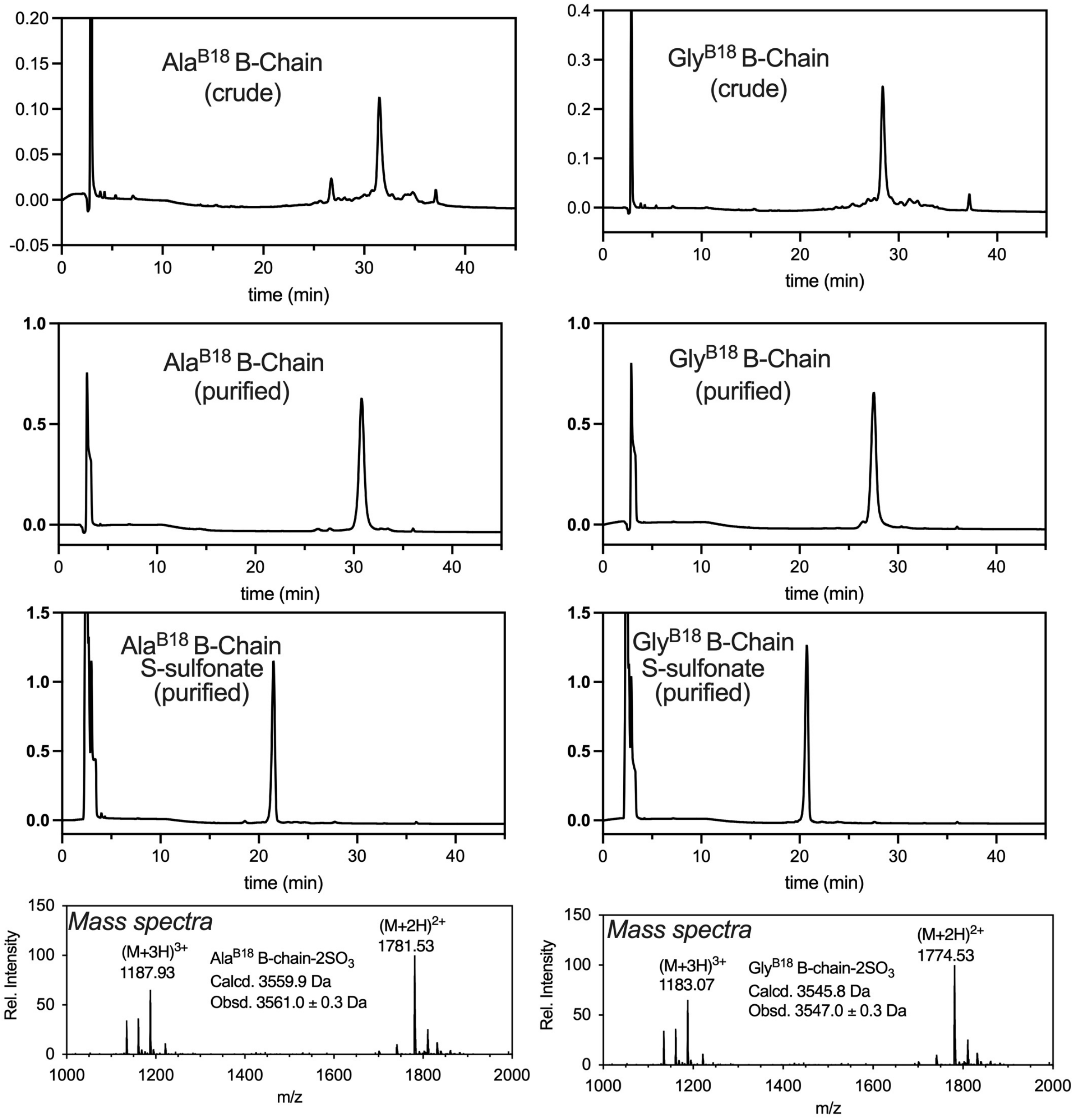

**Figure S12**. Synthesis, purification and S-sulfonation of variant $Ala^{B18}$ and $Gly^{B18}$ B-chain segments for chain combination reaction. LCMS analysis of S-sulfonates are provided at the bottom.

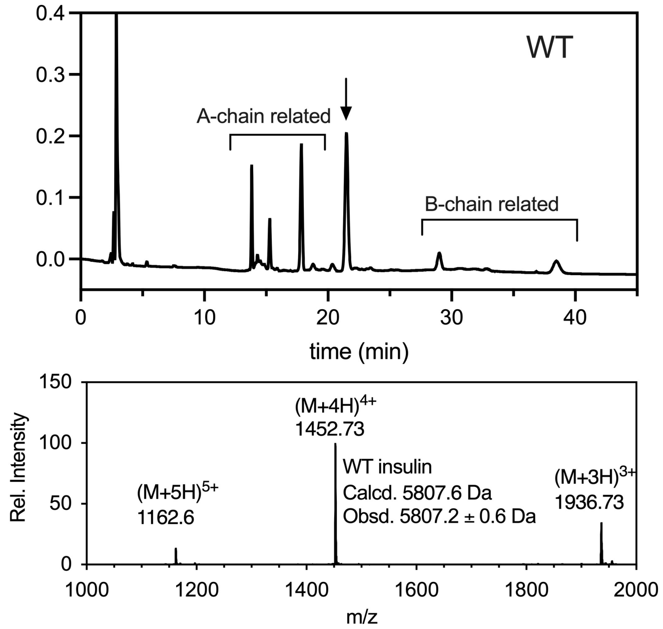

**Figure S13**. LC-MS analysis for the chain combination reaction of WT A-chain $SO_3$ and WT B-chain $SO_3$ in presence of DTT in 0.1 M glycine buffer at pH 10.5.

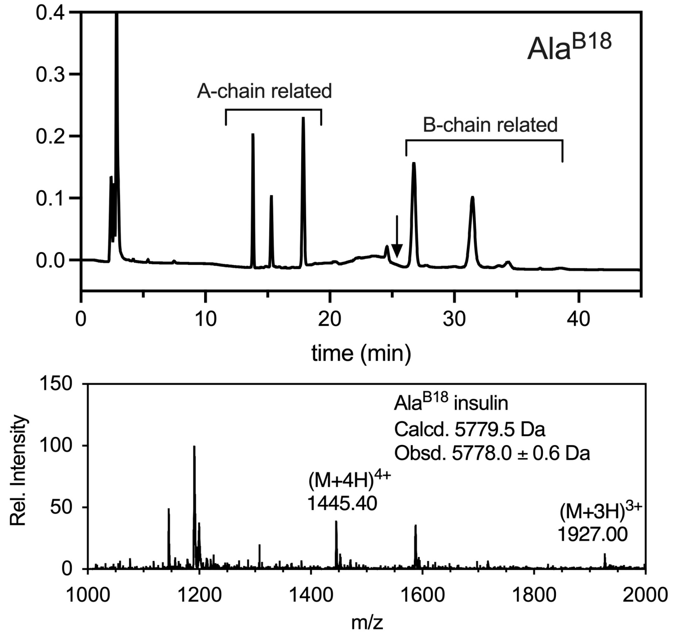

**Figure S14**. LC-MS analysis for the chain combination reaction of WT A-chain $SO_3$ and $Ala^{B18}$ B-chain $SO_3$ in presence of DTT in 0.1 M glycine buffer at pH 10.5.

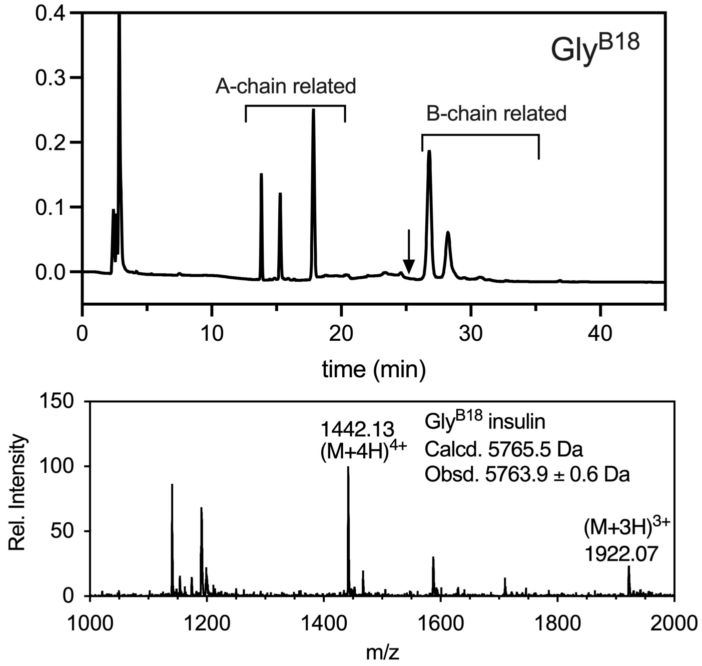

**Figure S15**. LC-MS analysis for the chain combination reaction of WT A-chain $SO_3$ and Gly$^{B18}$ B-chain $SO_3$ in presence of DTT in 0.1 M glycine buffer at pH 10.5.

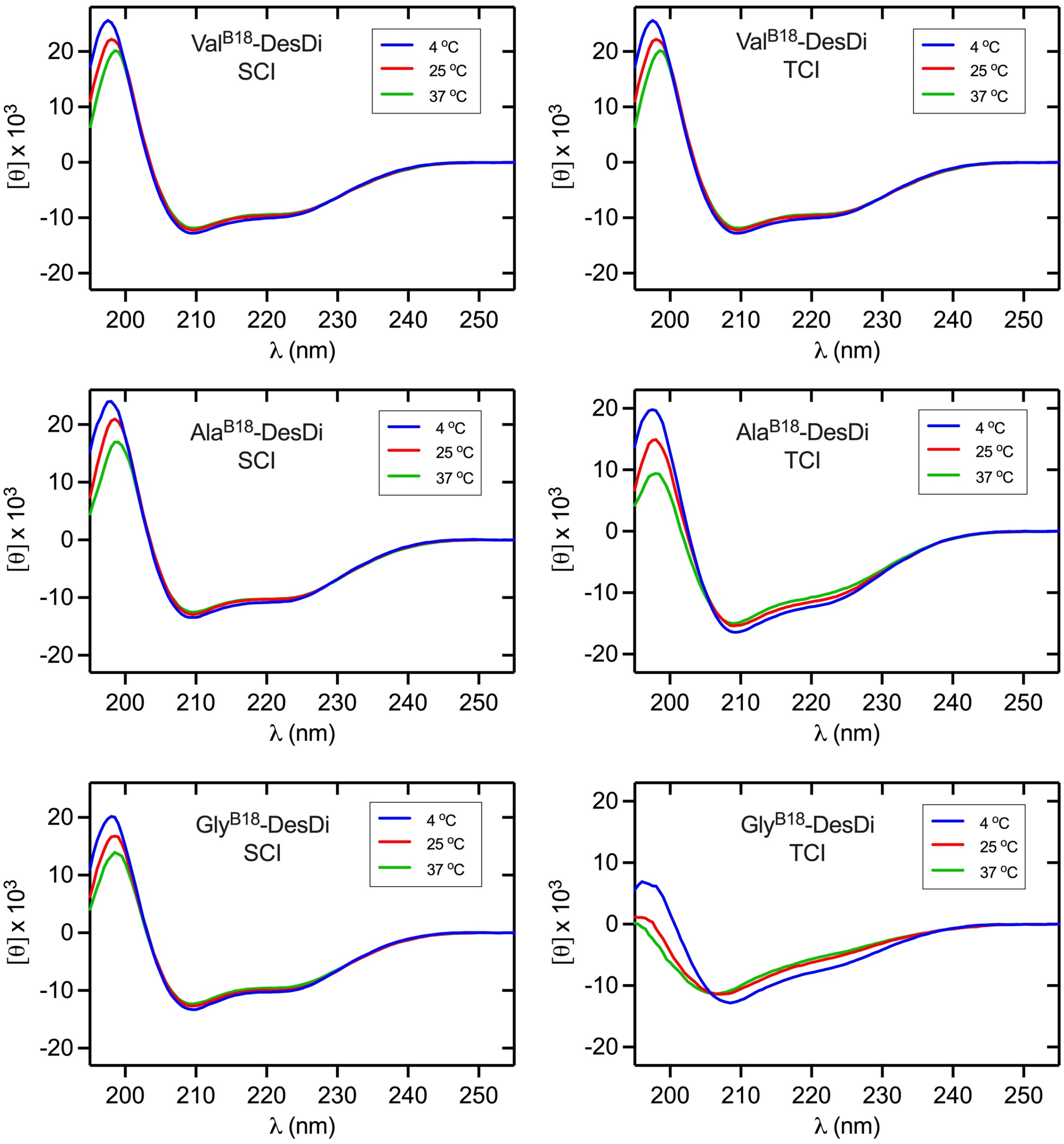

**Figure S16**. Far-UV CD spectra of DesDi single-chain and two-chain insulin analogs. The color code is inset within each panel.

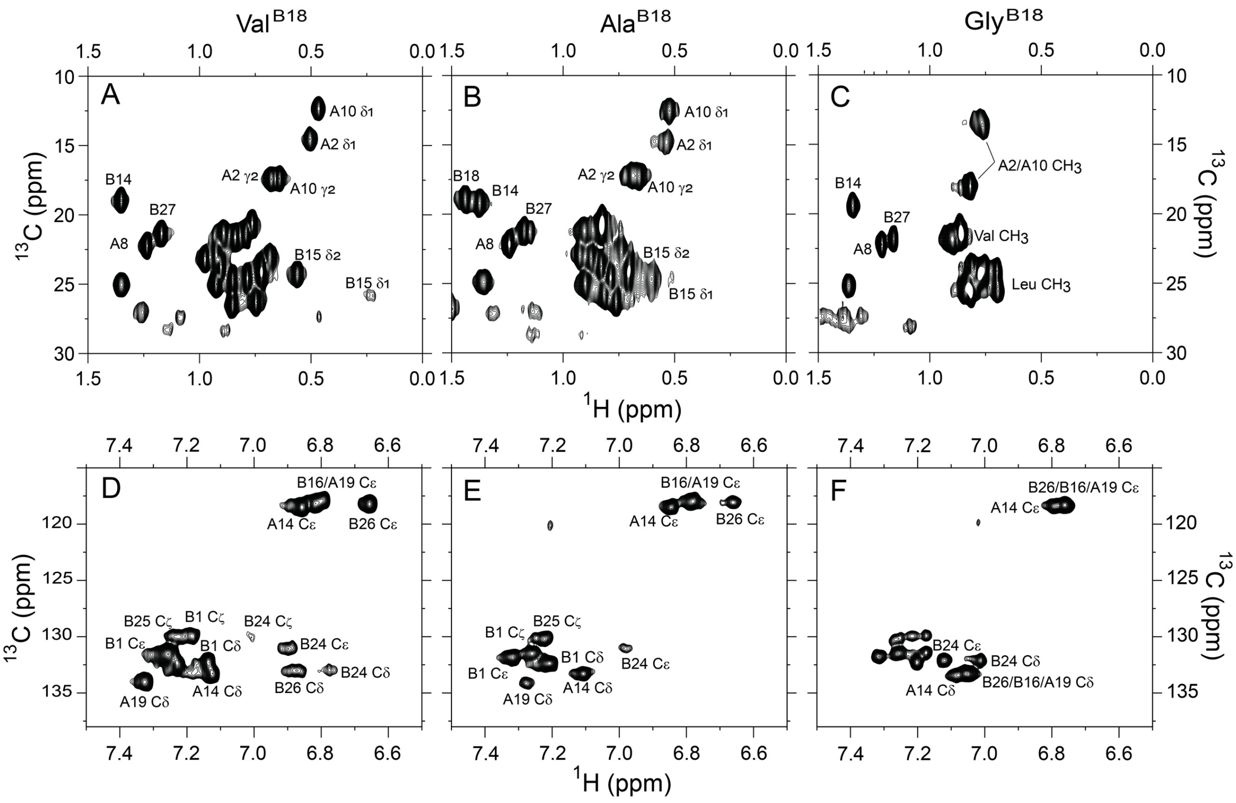

**Figure S17**. Nature abundance $^{1}$H-$^{13}$C HSQC NMR spectra of two-chain DesDi insulin analogs. Top panel provides the methyl region of $^{1}$H-$^{13}$C HSQC correlation spectra of (*A*) Val$^{B18}$ analog, (*B*) Ala$^{B18}$ analog and (*C*) Gly$^{B18}$ analog. Bottom panel is aromatic region of $^{1}$H-$^{13}$C HSQC spectra of (*D*) Val$^{B18}$ analog, (*E*) Ala$^{B18}$ analog and (*F*) Gly$^{B18}$ analog. Spectra were acquired at a $^{1}$H frequency of 700 MHz at pD 7.4 (direct meter reading) and 32 $^{o}$C. Selected resonance assignments are as indicated. HSQC, heteronuclear single-quantum coherence.

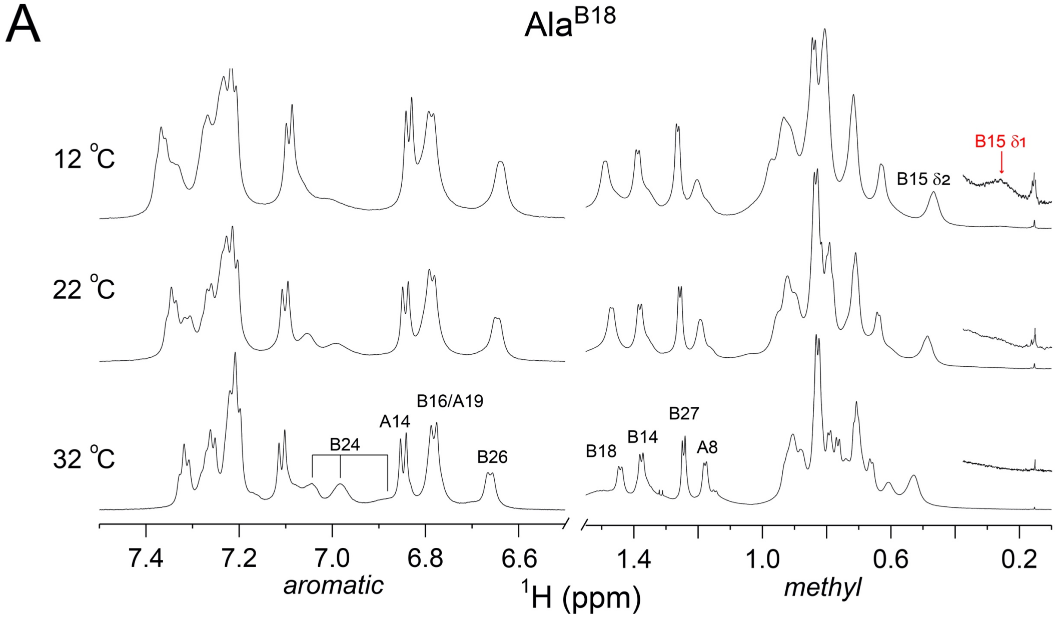

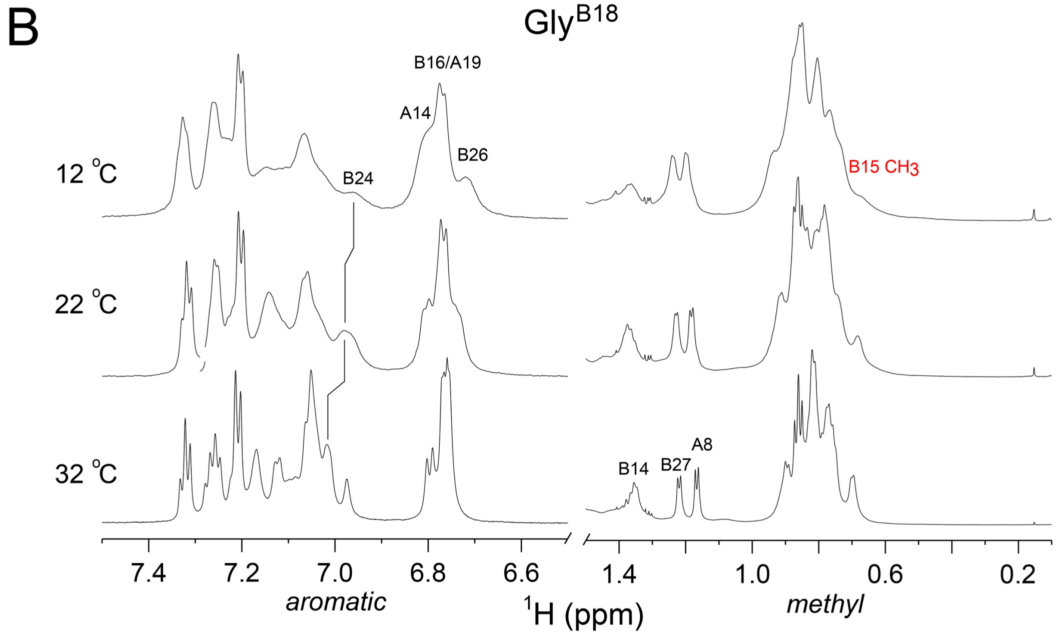

**Figure S18**. Temperature-dependent $^{1}$H-NMR studies. (*A*) 1D $^{1}$H-NMR spectra of two-chain DesDi-Ala$^{B18}$-insulin in aromatic and methyl region at 32 °C (*bottom*), 22 °C (*middle*) and 12 °C (*top*). 10 times intensity increase in the extended region (*inset* at far right). (*B*) $^{1}$H-NMR spectra of DesDi-Gly$^{B18}$-insulin in aromatic and methyl region at 32 °C (*bottom*), 22 °C (*middle*) and 12 °C (*top*). Spectra were acquired at a $^{1}$H frequency of 700 MHz at pD 7.4 (direct meter reading) and selected resonance assignments are as indicated.

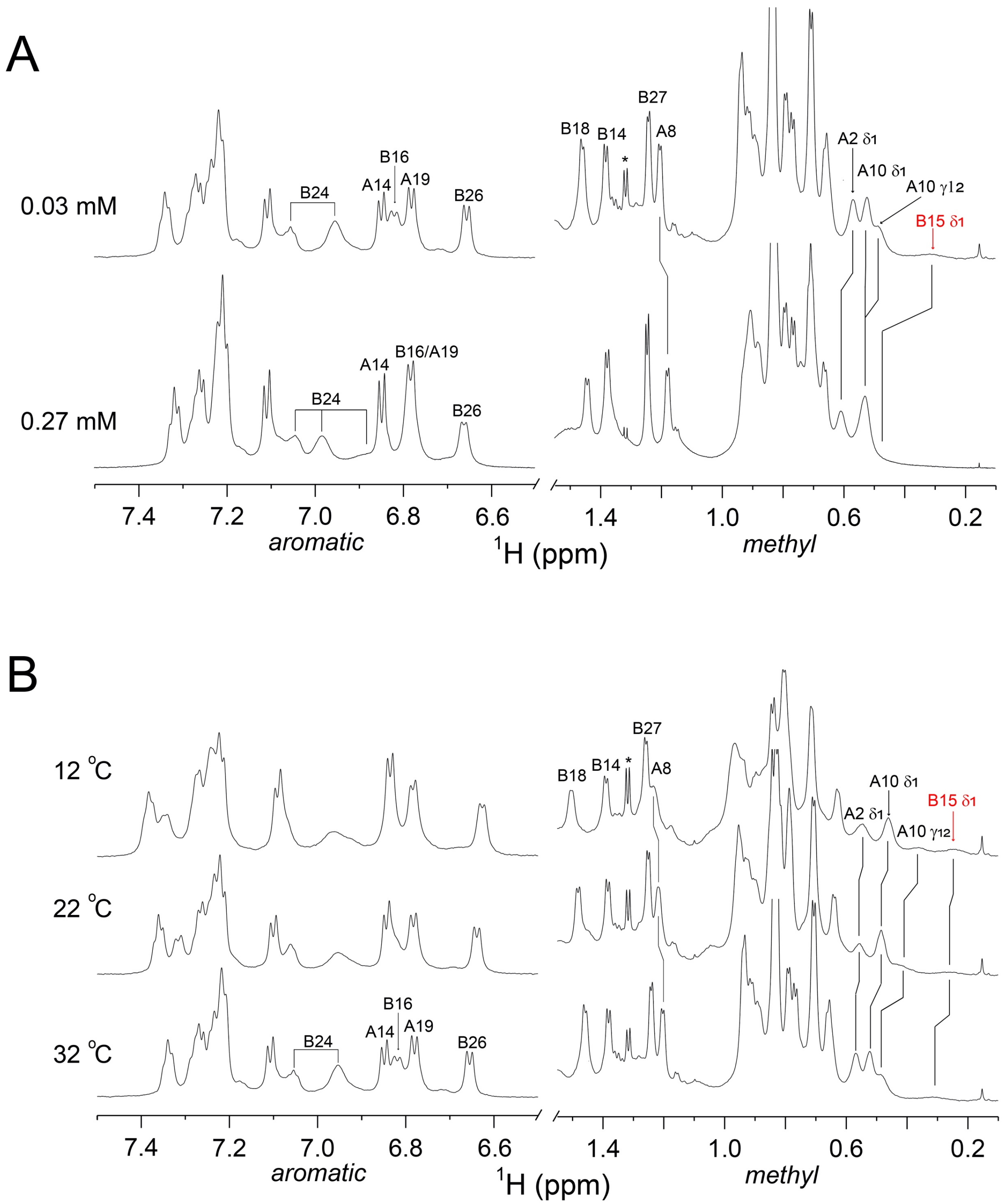

**Figure S19**. Concentration- and temperature-dependent NMR of two-chain DesDi-Ala$^{B18}$. (*A*) $^{1}$H-NMR spectra of DesDi-Ala$^{B18}$ in aromatic and methyl region at 32 ºC and pD 7.4 (direct meter reading) in 0.27 mM of concentration (*bottom*) and around 9 times dilution (0.03 mM; *top*). (*B*) $^{1}$H-NMR spectra of diluted two-chain DesDi-Ala$^{B18}$-insulin (0.03 mM) in aromatic and methyl region at 32 ºC (*bottom*), 22 ºC (*middle*) and 12 ºC (*top*). Spectra were acquired at a $^{1}$H frequency of 700 MHz at pD 7.4 (direct meter reading). Selected resonance assignments are as indicated; asterisk near 1.3 ppm (top) indicates contaminant.

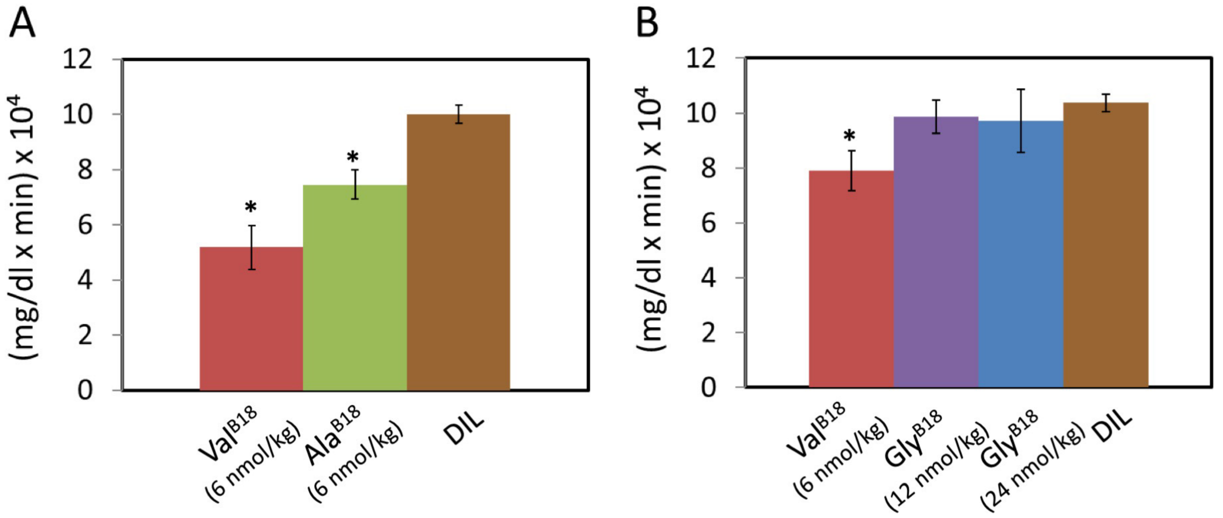

**Figure S20.** Analysis of STZ rat data. Areas under the curve (AUCs) pertaining to time course of blood-glucose concentrations for the subcutaneous (SQ) experiments is shown. The studies employed two-chain DesDi insulin and its B18 analogs. (A) $Val^{B18}$ (WT) DesDi and $Ala^{B18}$ at 6 nmol/kg rat; diluent at 100 μmol/kg rat (all at n =6). The AUC's of these two insulins are significantly different (indicated by asterisk) than diluent-injected control rats ($p$ value $< 0.05$). (B) $Val^{B18}$ (WT) DesDi (6 nmol/kg rat) compared with $Gly^{B18}$ (12 and 24 nmol/kg rat). $Val^{B18}$ is significantly different than dilute (indicated by asterisk) and p value for the $Gly^{B18}$ 12 nmol/kg rat dose is 0.07 and for the 24 nmol/kg rat dose is 0.22 with respect to $Val^{B18}$. There is no difference between the AUC's of $Gly^{B18}$ insulin analog relative to diluent control ($p$ values $>0.1$).

**Table S1**. Effects of B18 Substitutions (11,12) on Foldability[a,b]

| analog | Isolated yield (from 100 mg crude) |
|---|---|
| $Val^{B18}$ (WT) DesDi | 21.7 mg |
| $Ala^{B18}$ DesDi | 12.5 mg |
| $Gly^{B18}$ DesDi | 3.2 mg |

[a]Isolated yields of *in vitro* folding of single-chain insulin analogs in DesDi template ($Lys^{B28}$-des-dipeptide[B28. B29]-mini-proinsulin; "WT").

[b]Although the folding reactions employed crude synthetic products, the three polypeptides were prepared using the same protocol, suite of reagents and instrument. Analytical reverse-phase HPLC and LC/MS characterization of the crude products indicated comparable quality of solid-phase synthesis.

**Table S2**. CD derived helical parameters.

A. *Molar ellipticities at 222 nm and 208 nm at 25°C and pH 7.4*

| *analogs* | $[\theta]_{222} \times 10^3$ | $[\theta]_{208} \times 10^3$ |
|---|---|---|
| WT-DesDi-SCI[b] | -9.55 | -11.56 |
| $Ala^{B18}$-DesDi-SCI | -10.26 | -12.15 |
| $Gly^{B18}$-DesDi-SCI | -9.93 | -12.11 |
| WT DesDi-TCI[c] | -10.27 | -15.00 |
| $Ala^{B18}$-DesDi-TCI | -11.05 | -14.94 |
| $Gly^{B18}$-DesDi-TCI | -5.69 | -11.33 |

B. *Deconvolution estimates for helix, sheet and random-coil contents*[a]

| *analogs* | α-helix (%) | β-sheet (%) | Disordered (%) | Turn (%) |
|---|---|---|---|---|
| WT-DesDi-SCI[b] | 46.8 | 14.9 | 25.2 | 9.6 |
| $Ala^{B18}$-DesDi-SCI | 50.0 | 13.4 | 25.2 | 9.3 |
| $Gly^{B18}$-DesDi-SCI | 46.2 | 14.2 | 27.9 | 9.5 |
| WT DesDi-TCI[c] | 45.2 | 14.6 | 28.4 | 13.5 |
| $Ala^{B18}$-DesDi-TCI | 47.5 | 13.6 | 26.1 | 14.1 |
| $Gly^{B18}$-DesDi-TCI | 20.6 | 25.6 | 30.8 | 20.5 |

[a]Total percent α-helix, β-sheet, and disordered coil were obtained from spectra acquired at 25 °C using the SELCON-3 algorithm.

[b]SCI designated single-chain insulin, the present synthetic precursor (49 residues; $Lys^{B28}$-*des*-dipeptide[B29, B30]-mini-proinsulin).

[c]TCI designates two-chain insulin, obtained from DesDi SCI precursor by Endo-Lys-C endoprotease cleavage (see Experimental Procedures).

**Table S3**. Responses of ER stress markers to mutations $Ala^{B18}$ and $Gly^{B18}$ in human proinsulin (3,13)[a]

| | relative activity ratio in Western blot[b] (normalized by GAPDH) | | | relative activity ratio in real-time-q-PCR (mRNA abundances)[c] | |
|---|---|---|---|---|---|
| | pPERK/PERK | CHOP | BiP | *CHOP* | *BiP* |
| $Val^{B18}$ | 1.00±0.01 | 1.00±0.01 | 1.00±0.01 | 1.00±0.05 | 1.00±0.1 |
| $Ala^{B18}$ | 35.5±1.03 | 4.6±0.51 | 12.1±0.39 | 1.96±0.16 | 5.6±1.68 |
| $Gly^{B18}$ | 65.1±3.05 | 7.3±2.54 | 37.3±1.96 | 4.42±0.75 | 10.56±2.11 |

[a]The proinsulin constructs employed in transient transfection also contained an N-terminal Myc epitope tag.
[b]Values for B18 mutants were independently normalized in each blot with respect to $Val^{B18}$ (WT), defined as 1.00. Values of pPERK/PERK thus provide a ratio of ratios (5).
[c]Values for B18 mutants were independently normalized for each messenger RNA with respect to $Val^{B18}$ (WT), defined as 1.00.

**Table S4**. B18 Methyl-Related Distances[a,b]

| | $Val^{B18}$ | | $Ala^{B18}$ |
|---|---|---|---|
| | γ1 | γ2 | |
| $Leu^{A13}$ δ1 | | 4.5 | 5.9 |
| $Leu^{A13}$ δ2 | | 4.0 | 5.5 |
| $Leu^{A16}$ δ1 | | 4.4 | 6.9 |
| $Leu^{A16}$ δ2 | | 6.3 | 5.3 |
| $Leu^{B15}$ δ1 | | | 5.6 |
| $Leu^{B15}$ δ2 | 5.9 | | 7.8 |
| $Leu^{B15}$ δ2 | | 6.0 | |
| $Glu^{A17}$ δ | 4.7 | | 6.3 |
| $Cys^{B19}$ SG | 4.4 | | 6.0 |

[a]Side-chain distances (Å) from the methyl carbons of $Val^{B18}$ and $Ala^{B18}$ to neighboring residues in the native state as observed in a crystallographic protomer or as predicted by a rigid-body model.

[b]Distances in WT insulin (Å) were measured in an insulin monomer (extracted from 2-Zn insulin; PDB entry 4INS (10)) using PyMOL.
